# Syndecan-1 Promotes Alveolar Type 2 Epithelial Cell Senescence during Lung Fibrosis

**DOI:** 10.64898/2026.03.16.712248

**Authors:** Changfu Yao, Milena Espinola, Xue Liu, Yizhi Wang, Marilia Zuttion, Virinchi Kuchibhotla, Xuexi Zhang, Larissa Langhi Prata, Silvia Cho, Zackery Ortega, Emily Braghramian, Kimberly Merene, Ying Wang, Susan Jackman, Antonina Caudill, Fatima Contreras, Jiurong Liang, Dianhua Jiang, Paul W Noble, Cory M. Hogaboam, Barry R. Stripp, Cecilia Lopez-Martinez, Sina A. Gharib, Amara Seng, Nunzio Bottini, William C. Parks, Peter Chen, Tanyalak Parimon

## Abstract

Idiopathic pulmonary fibrosis (IPF) is an age-related, progressive, and fatal interstitial lung disease for which effective therapies remain limited. Alveolar type 2 (AT2) epithelial cells serve as facultative stem cells essential for alveolar repair; however, AT2 cell senescence disrupts epithelial regeneration and contributes to fibrotic remodeling in IPF. Syndecan-1 is a transmembrane heparan sulfate proteoglycan predominantly expressed by lung epithelial cells, but its role in AT2 dysfunction during fibrosis is poorly defined. Here, we demonstrate that syndecan-1 is robustly upregulated in AT2 cells in IPF and other fibrotic lung diseases, as well as in murine bleomycin-induced lung fibrosis. Syndecan-1 expression was further enhanced with aging and associated with increased fibrotic burden in aged mice. Using integrated human transcriptomic analyses, mouse genetic models, and epithelial cell–based systems, we show that excess syndecan-1 promotes cell-autonomous epithelial senescence and impairs AT2 progenitor function. Elevated syndecan-1 reduced AT2 renewal capacity, disrupted differentiation, and diminished surfactant protein C level, whereas genetic loss of syndecan-1 attenuated senescence and preserved epithelial function following injury. Together, these findings identify syndecan-1 as a critical epithelial regulator of AT2 senescence and maladaptive repair in pulmonary fibrosis and support targeting syndecan-1–driven epithelial dysfunction as a potential therapeutic strategy.

## Introduction

Chronic respiratory diseases, including interstitial lung diseases, are a significant cause of morbidity and mortality worldwide (1). Idiopathic pulmonary fibrosis (IPF) is a progressive, fatal disorder with rising global incidence and no curative therapy (2). Beyond lung transplantation, no treatments effectively halt or reverse the pathological fibroproliferative remodeling that defines this disease. Thus, identifying the key molecular drivers that underlie fibrosis initiation and progression is essential for developing targeted and effective interventions.

Alveolar type 2 (AT2) epithelial cells function as the lung progenitor stem cells (3), proliferating and differentiating into alveolar type 1 (AT1) cells to maintain alveolar homeostasis and mediate effective repair following injuries (4–8). When AT2 function is impaired, dysplastic alveolar repair ensues, driving the initiation and progression of lung fibrosis (4, 6, 9). Converging experimental and compelling evidence implicates cellular senescence as a central driver of AT2 dysfunction and exhaustion, directly contributing to dysplastic repair and lung fibrosis (10–17). While multiple molecules that promote AT2 senescence have been described (14, 16, 18–21), relatively few upstream molecules governing this process have been defined (22, 23).

Syndecan-1 is a type I transmembrane heparan sulfate proteoglycan expressed on the basolateral surface of airway and alveolar epithelial cells, where it regulates multiple cellular processes through interactions between its heparan sulfate chains and soluble or matrix-associated ligands (24). During normal epithelial repair, shedding of syndecan-1 relieves this regulatory constraint, thereby promoting epithelial differentiation (25, 26) and re-epithelialization (27–29). Syndecan-1 is upregulated in AT2 cells and exhibits profibrotic activity in mice by suppressing miRNAs that target the cellular senescence pathway (30). In addition, syndecan-1 attenuates epithelial apoptosis during influenza infection (31), a feature consistent with the apoptosis-resistant phenotype of senescent cells. Together, these findings suggest that excess syndecan-1 in AT2 cells promotes AT2 senescence in a cell–autonomous manner and contributes to maladaptive alveolar repair in pulmonary fibrosis.

In this study, we identify syndecan-1 as a regulator of AT2 cell senescence using complementary human lung tissues and murine models. We show that excess syndecan-1 expression in AT2 cells promotes AT2 senescence and impairs their reparative capacity. These defects are associated with maladaptive alveolar repair in fibrotic lung injury. Mechanistically, we find that syndecan-1–mediated AT2 senescence is linked to increased p53 acetylation. Our findings establish excess epithelial syndecan-1 as a key determinant of AT2 cell senescence, leading to defective alveolar repair and fibrotic lung remodeling.

## Results

### Syndecan-1 is upregulated in AT2 cells in idiopathic pulmonary fibrosis and other fibrotic lung diseases

We previously reported that syndecan-1 is upregulated in lung epithelial cells across interstitial lung diseases (ILDs), including IPF (30). In the current study, we focused specifically on alveolar type 2 (AT2) cells. Two independent analyses– one using a published IPF and control lung single-cell RNA sequencing (scRNA-seq) dataset (GSE122960) (16) (**Fig. 1A**), and another using an unpublished integrative dataset comprising 80 control and 85 IPF lung scRNA-seq samples **(Supplementary Fig. S1A**)– demonstrated a significant increase in syndecan-1 (SDC1) mRNA expression in AT2 cells in IPF compared with controls. Consistent with these findings, spatial transcriptomic analysis of control (N = 3) and IPF (N = 2) lungs (GSE292956) revealed an upregulation (**Fig. 1 B–C)** and co-expression of *SDC1* with *ABCA3,* a canonical marker of AT2 cells (**Fig. 1 D).** Immunofluorescence analysis of explant lung tissues confirmed a significant increase in SDC1 protein abundance in AT2 cells in IPF lungs, compared with the controls (**Fig. 1E**). Notably, syndecan-1 expression was also upregulated in other fibrotic ILDs, including rheumatoid arthritis–associated ILD (**Supplementary Fig. S1B**), supporting a broader role for syndecan-1 across the spectrum of fibrotic lung diseases. Collectively, these findings establish syndecan-1 as a conserved molecular feature of fibrotic AT2 cells, prompting us to next assess the translational consequences of excess syndecan-1 on AT2 cell function.

**Figure 1.**
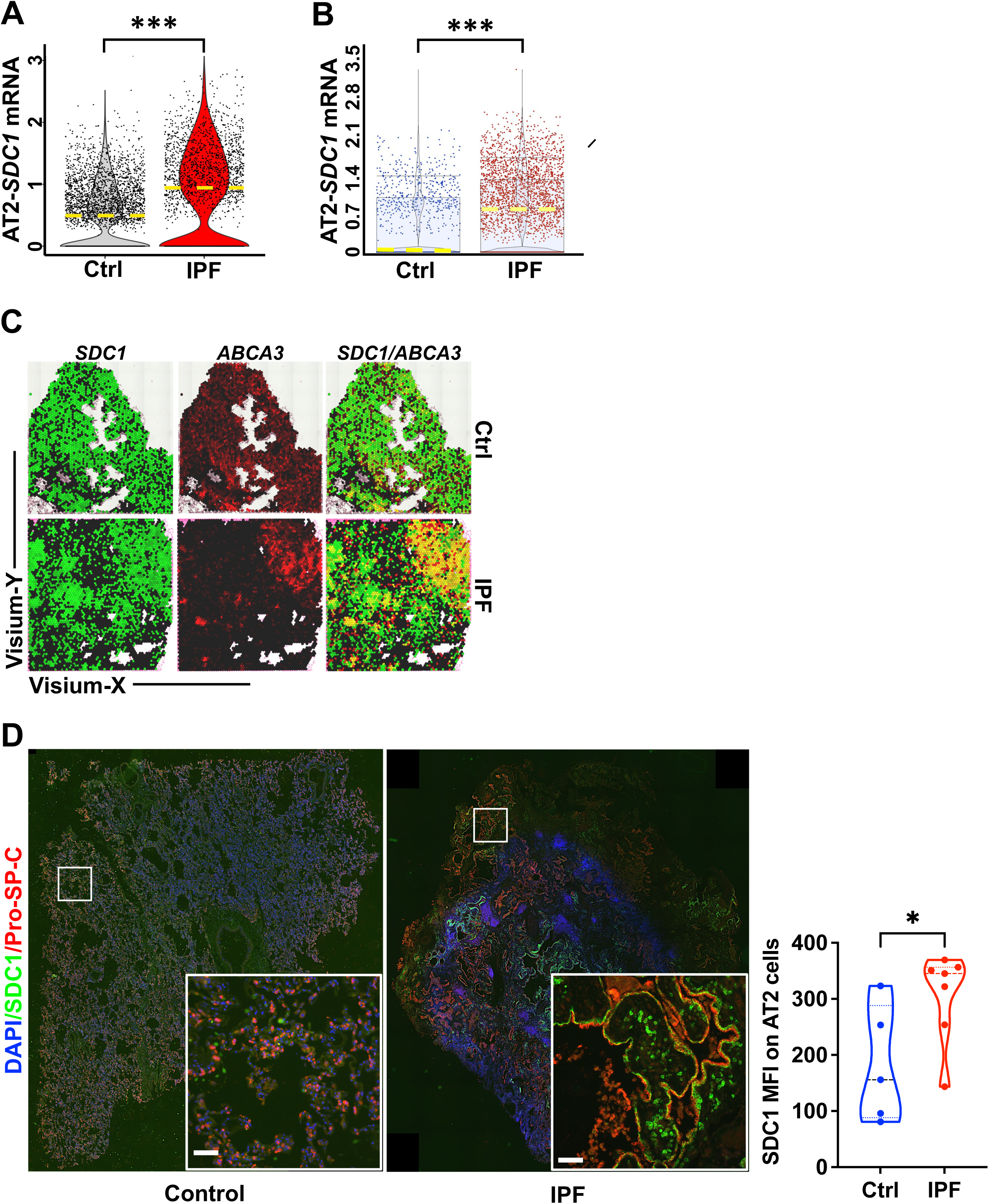
Syndecan-1 is overexpressed by alveolar type 2 (AT2) cells in Idiopathic Pulmonary Fibrosis (IPF). **A** Violin plot showing syndecan-1 (SDC1) mRNA expression level on AT2 cells from a scRNAseq dataset **(GSE122960)** comparing control (N=7) and IPF (N=9). **B–C** Analysis of a spatial transcriptomic cohort (**GSE292956**) from control (N=3) and IPF (N=2) lungs. **(B)** Violin plots depicting syndecan-1 expression on AT2 cells from controls and IPF lungs. **(C)** Spatial distribution of syndecan-1 (*SDC1*) and the AT2 cells (*ABCA3*) in control and IPF lungs. **D** Representative immunofluorescence images of control and IPF explant lung sections stained for syndecan-1 (SDC1; Alexa Fluor 488, green) and AT2 cells (pro–SP-C; Alexa Fluor 647, red). Syndecan-1 mean fluorescence intensity (MFI) was quantified in pro–SP-C⁺ AT2 cells across whole-lung sections from 5 controls and 7 IPF subjects. Scale=100 µm. *FDR<0.01 was considered statistically significant. *p<0.05; **p<0.005; ***p<0.0005 by Kruskal−Wallis, one-way ANOVA analysis, and Student’s t-test for two-sample comparison.

***Syndecan-1 is upregulated in aged AT2 cells and promotes pulmonary fibrosis in aged mice.*** To define the age-dependent regulation of syndecan-1 in fibrotic lung injury, we analyzed Sdc1 mRNA expression in AT2 cells from uninjured and bleomycin-injured lungs of young and aged mice using a published scRNA-seq dataset (GSE186246). An age-dependent increase in Sdc1 expression was observed specifically in Day 30 bleomycin-injured AT2 cells, whereas no differences were detected in uninjured AT2 cells (**Fig. 2A**). Consistent with these transcriptional findings, syndecan-1 protein levels were increased in AT2 cells from both young and aged mice following 21 days of bleomycin administration (**Fig. 2B and Supplementary Fig. S2A–B and S3A–B**), while no age-dependent differences were observed under uninjured conditions (**Supplementary Fig. S4A–B, S5A–B, and S6A–B**). Notably, image quantification of whole-lung sections showed that syndecan-1 expression was highest in aged, day 21, bleomycin-injured AT2 cells, but this increase did not reach statistical significance compared with the matched young mice (**Fig. 2C**).

**Figure 2.**
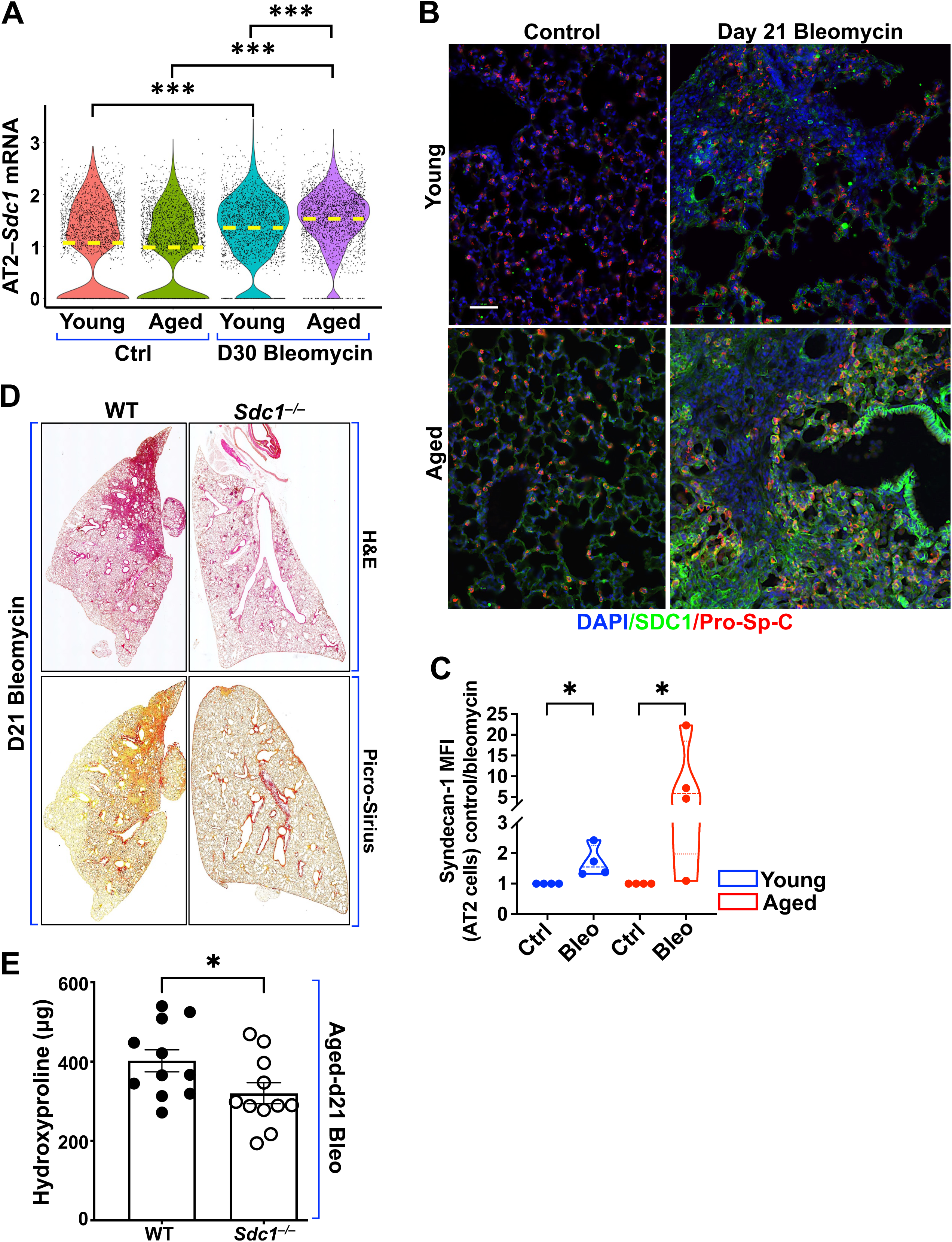
Overexpression of syndecan-1 by alveolar type 2 (AT2) cells in aged bleomycin–injured mice promotes lung fibrosis. **A** Violin plots of syndecan-1 expression in AT2 cells from uninjured and bleomycin-injured lungs of young and aged mice, based on the publicly available scRNA-seq dataset (**GSE186246**). **B** Representative immunofluorescence images of lung sections from control (uninjured) and day 21 bleomycin-injured young and aged mice, stained for syndecan-1 (Sdc1; Alexa Fluor 488, green), and AT2 cells (pro-SP-C; Alexa Fluor 546, red. Scale=50 µm. **C** Syndecan-1 mean fluorescence intensity (MFI) in pro–SP-C⁺ AT2 cells was quantified on whole lung sections of bleomycin-injured and uninjured lungs (**Supplemental Fig. S4, S5A–B, and S6A–B**). Analyses were performed on 4 mice per group using the Nikon GA3 image analysis software. **D–E** Representative images of H&E and Picro-Sirius staining **(D)**. Quantification of hydroxyproline content in the right lungs of aged wild-type (WT; N=11) and *Sdc1^–/–^*(N=11) mice following day 21 bleomycin-induced lung fibrosis (E). *FDR<0.01 was considered statistically significant. *p<0.05; **p<0.005; ***p<0.0005 by Kruskal−Wallis, one-way ANOVA analysis, and Student’s t-test for two-sample comparison.

To determine the functional consequences of excess syndecan-1 in AT2 cells in vivo, we utilized the bleomycin-induced lung injury and fibrosis model to assess fibrotic burden. Aged bleomycin-injured wild-type (WT) mice exhibited significantly greater collagen deposition per lung compared with age-matched *Sdc1^–/–^* mice, as measured by hydroxyproline lung content (**Fig. 2D**), a finding corroborated by H&E and Pico Sirius staining (**Fig. 2E**). Together, these data indicate that elevated syndecan-1 expression exacerbates fibrotic remodeling in the aged lung. Given the well-established relationship between aging and cellular senescence, and our prior findings that syndecan-1 engages senescence-associated pathways to exacerbate lung fibrosis in young mice, we next tested the hypothesis that syndecan-1 directly promotes AT2 cell senescence, thereby driving maladaptive repair and fibrotic progression in pulmonary fibrosis.

### Upregulation of syndecan-1 in AT2 cells is associated with AT2 senescence

Given the well-established relationship between aging and cellular senescence, and our prior findings that syndecan-1 engages senescence-associated pathways to exacerbate lung fibrosis in young mice (30), we next tested the hypothesis that syndecan-1 directly promotes AT2 cell senescence, thereby driving maladaptive repair and fibrotic progression in pulmonary fibrosis. To further define the relationship between syndecan-1 and AT2 cell senescence, we first assessed the composite transcriptomic senescence score (SenMayo Senescence Score; **Supplementary Table 1**) (32) in AT2 cells from IPF and control lungs using a published scRNA-seq dataset (GSE122960). IPF AT2 cells exhibited significantly higher SenMayo senescence scores compared with control AT2 cells (**Fig. 3A and Supplementary Fig. S7A**), a finding that was independently recapitulated in a spatial transcriptomics dataset (GSE292956, **Supplementary Fig. S7B**). Stratification of AT2 cells by syndecan-1 expression level—high (*SDC1*^Hi^) versus low (*SDC1*^Low^)—revealed a positive association between SDC1 expression and senescence score (**Fig. 3B**). To validate these transcriptomic findings at the protein level, we immunostained control and IPF lung explants for the senescence marker p21 and quantified the proportion of p21⁺ cells within the SDC1⁺ AT2 cell population. IPF lungs demonstrated a significantly higher percentage of p21⁺SDC1⁺ double-positive cells compared with controls (**Fig. 3C and Supplementary Fig. S7C**), supporting a link between elevated syndecan-1 expression and AT2 cell senescence in human fibrotic lungs.

**Figure 3.**
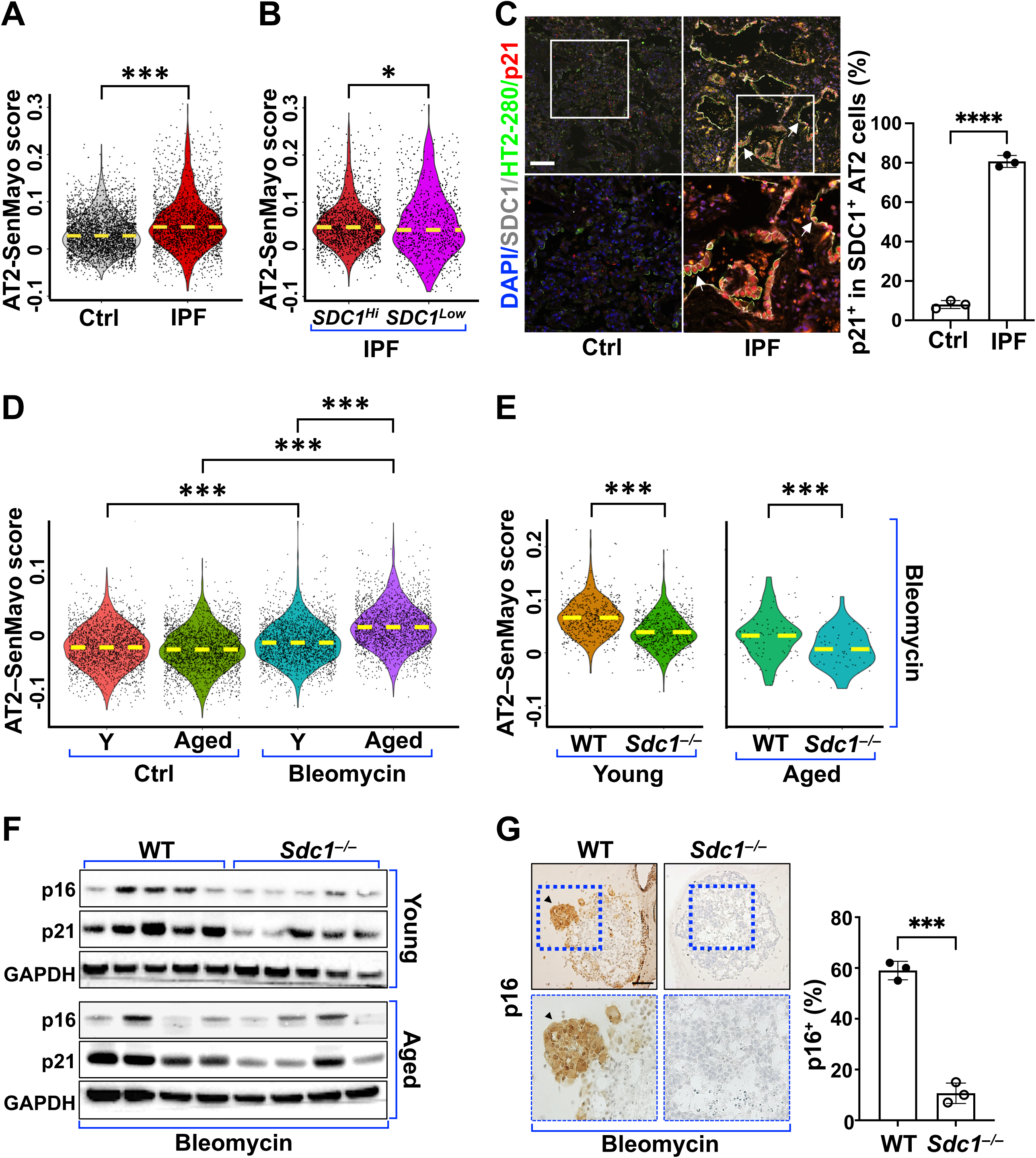
Upregulation of syndecan-1 in alveolar type 2 epithelial cells (AT2) promotes AT2 cellular senescence during lung fibrosis. **A–B** Violin plots showing the mean composite transcriptomic senescence score (SenMayo Senescence Score) in AT2 cells from control and IPF lungs based on the scRNA-seq dataset (**GSE122960**). **(A)** Senescence scores across all AT2 cells. (**B)** Comparison of senescence scores between high syndecan-1–expressing (*SDC1^Hi^*) and low syndecan-1–expressing (*SDC1*^low^) AT2 cells within each group. **C** Representative immunofluorescence images of control (N=3) and IPF (N=3) lung sections stained for syndecan-1 (SDC1; Alexa Fluor 488, green) and p21 (Alexa Fluor 647, white). Quantification of % p21^+^ on SDC1^+^ cells were performed in the lung sections of 3 controls and 3 IPF lungs. Scale bar=100 μm. **D–E** SenMayo Senescence Score of mouse AT2 cells: **(D)** AT2 cells from young and aged WT mice under control and bleomycin-injured conditions (**GSE186246**). **(E)** AT2 cells from young (**GSE134948**) and aged (**GSE292961**) WT *and Sdc1^–/–^*mice following bleomycin-induced lung fibrosis at days 21 and 28, respectively. **F** Western blot immunostaining for p16 and p21 in whole lung homogenates from young and aged WT and *Sdc1^–/–^* mice following bleomycin injury, N=5 per group. **G** Representative immunohistochemistry images of bleomycin-stimulated (50 µg/mL via the lower chamber for 48 hours) AT2 alveolospheres derived from young WT and *Sdc1^–/–^* mice staining for p16, N=3 per group. The percentage of p16^+^ cells was quantified in each mouse AT2 alveolospheres. *\**FDR*<*0.01 is considered significant. *\**p<0.05*; \*\**p<0.005*; \*\*\**p<0.0005 by Kruskal−Wallis, one-way ANOVA analysis for multiple comparisons, and Student’s t-test for two-sample comparisons.

Complementing these human data, we analyzed AT2 cells from young and aged mice in a published scRNA-seq dataset (GSE186246) and found significantly higher SenMayo senescence scores in bleomycin-injured AT2 cells from both young and aged mice compared with uninjured controls (**Fig. 3D**). Notably, among bleomycin-injured mice, aged AT2 cells exhibited higher senescence scores than AT2 cells from young mice, whereas no age-dependent differences were observed under uninjured conditions (**Fig. 3D**). To directly assess the contribution of syndecan-1 to AT2 senescence, we compared AT2 cell transcriptomes from young (GSE134948) and aged (GSE292961) WT and *Sdc1^–/–^* mice (**Supplementary Fig. S8–S9**). Bleomycin-injured *Sdc1^–/–^* AT2 cells exhibited significantly lower SenMayo senescence scores compared with WT AT2 cells in both young and aged cohorts (**Fig. 3E**, **Supplementary Fig. S10A–B**). Consistently, expression of senescence markers p16 and p21 was reduced in young bleomycin-injured *Sdc1^–/–^* compared with WT whole-lung homogenates (**Fig. 3F and Supplementary Fig. S10C**). Collectively, these findings demonstrate a conserved association between syndecan-1 expression and AT2 cell senescence across human and murine systems, supporting syndecan-1 as a key regulator of epithelial senescence in fibrotic lung disease.

### Syndecan-1 mediates lung epithelial cell senescence

To implicate a direct role for syndecan-1 in AT2 cell senescence, we generated AT2 alveolospheres from FACS-sorted SftpcCreER.R26-tdTomato*^+^*cells of uninjured WT and *Sdc1^–/–^* mice and assessed senescence following bleomycin exposure. After 48 hours of bleomycin stimulation, WT AT2 alveolospheres exhibited a significantly higher proportion of p16⁺ cells compared with *Sdc1^–/–^* alveolospheres (**Fig. 3G**). Similar reductions in p16 and p21 expression were observed following doxorubicin-induced senescence (**Supplementary Fig. S10D**).

To further define the role of syndecan-1 in promoting lung epithelial cell senescence, we utilized both mouse and human lung epithelial cell models. Senescence was first assessed in mouse lung epithelial MLE-15 cells engineered to overexpress (OE) or delete (*Sdc1* knockout; KO) syndecan-1, with an empty vector or a non-targeting guide RNA serving as control (Ctrl). Syndecan-1 expression levels were confirmed by flow cytometry (**Supplementary Fig. S11A–S11D**). In unperturbed MLE-15 cells, overexpressing syndecan-1 in the cells (MLE-15-OE) without cell stimulation increased p16 and p21 expression higher than the controls (**Fig. 4A**). In contrast, MLE-15-KO h exhibited reduced p21 expression compared with control cells both at baseline and following bleomycin stimulation (**Fig. 4A**). Additionally, serum-starvation MLE-15 cells for 48 h increased senescence-associated β-galactosidase (SA-β-gal) activity in the OE cells the highest, followed by WT and KO cells, respectively (**Fig. 4B).** The data suggested that alteration of syndecan-1 positively correlated with cellular senescence. Consistent with these findings, silencing or knockdown of SDC1 in human lung epithelial cell lines (BEAS-2B and A549 cells (**Supplementary Fig. S11C–D**) resulted in reduced p21 expression and attenuated SA-β-gal activity relative to controls (**Fig. 4C–D**). Restoration of syndecan-1 expression in SDC1-silenced BEAS-2B cells reversed these effects (**Fig. 4C**). Collectively, these data indicate that syndecan-1 expression directly and positively regulates epithelial cell senescence in a cell-autonomous manner.

**Figure 4.**
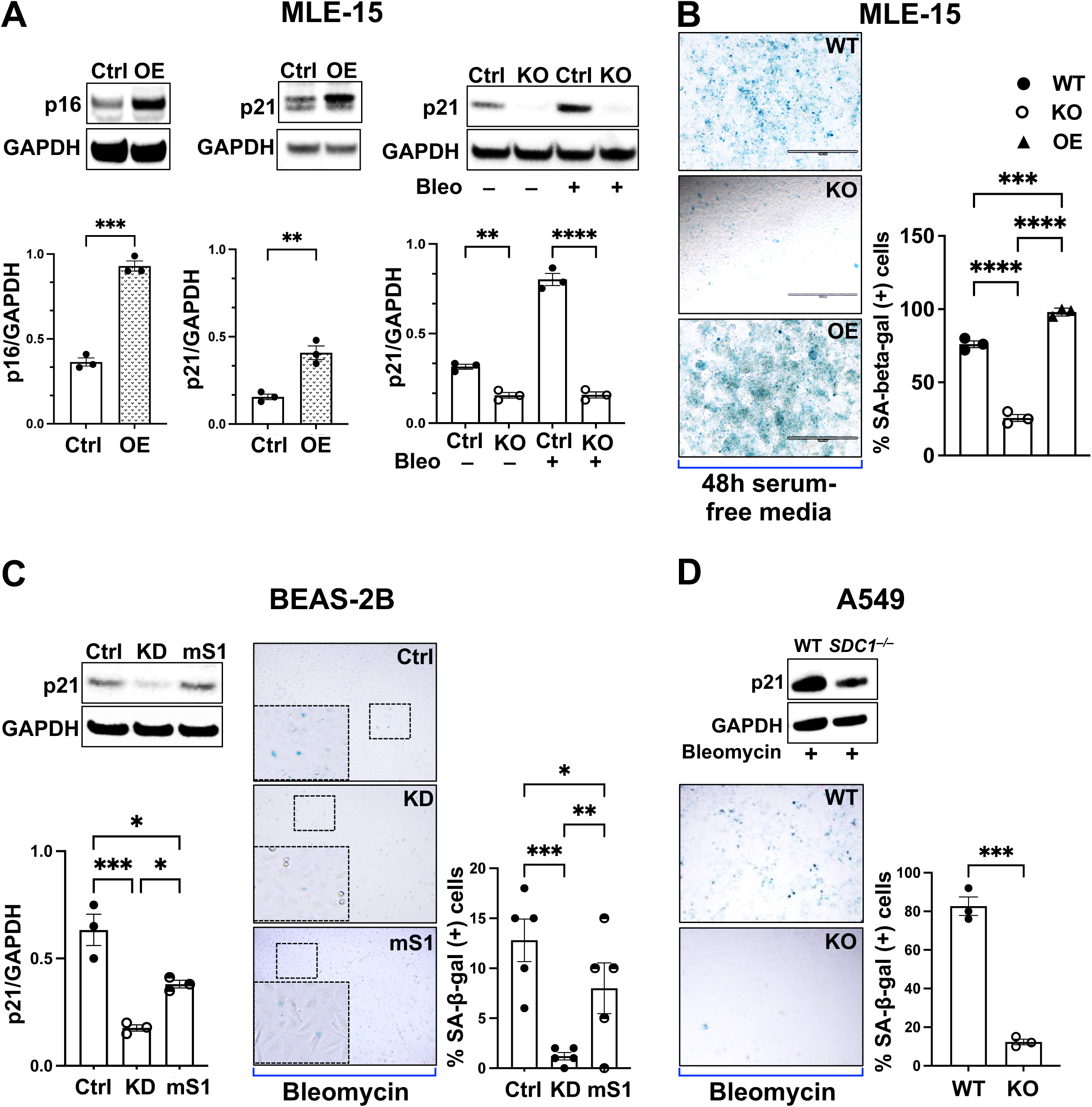
Overexpression of syndecan-1 is associated with the propensity of alveolar type 2 epithelial (AT2) senescence *in vitro*. **A–D** Western blotting (WB) for p16 and p21 and senescence-associated (SA) β-galactosidase activity of cell lysates from control and bleomycin–stimulated conditions in mouse and human lung epithelial cell types with different syndecan-1 expression levels. Densitometry quantification of p16 or p21 was normalized to GAPDH. **(A)** and **(B)** MLE-15 scramble control (Ctrl), overexpressed (OE), and CRISPR–deletion syndecan-1 (KO) for p16 and p21 and for SA-β-galactosidase activity, respectively. **(C)** BEAS-2B scramble control, syndecan-1 knockdown (KD), and mouse syndecan-1 were supplemented to the syndecan-1 knockdown cells (*KD+mS1*) for p21 WB and SA-β-galactosidase activity. **(D)** A549 scramble control and CRIPRS-deletion syndecan-1 (*SDC1^–/–^*) cells. *\**p<0.05*; \*\**p<0.005*; \*\*\**p<0.0005 by Kruskal−Wallis, one-way ANOVA analysis for multiple comparisons, and Student’s t-test for two-sample comparisons.

***Excess syndecan-1 impairs AT2 cell renewal and differentiation.*** We next examined the impact of excess syndecan-1 on AT2 cell functional capacity using AT2 alveolosphere assays. Under uninjured conditions, colony-forming efficiency (CFE) was comparable between WT and *Sdc1^–/–^* AT2 alveolospheres (**Fig. 5A**). Bleomycin injury significantly reduced CFE in both genotypes; however, the reduction was more pronounced in WT compared with *Sdc1^–/–^* alveolospheres (**Fig. 5A**). Assessment of AT2 differentiation revealed normal alveolosphere morphology in uninjured WT and *Sdc1^–/–^* cultures (**Supplementary Fig. S12A**). In contrast, bleomycin-injured WT AT2 alveolospheres exhibited aberrant morphology and dysregulated AT1 marker (Pdpn) localization, suggestive of impaired differentiation (**Fig. 5B, Supplementary Fig. S12B–C**). These abnormalities were attenuated in bleomycin-injured *Sdc1^–/–^* alveolospheres. Together, these findings suggest that elevated syndecan-1 levels compromise AT2 renewal and disrupt normal differentiation following lung injury.

**Figure 5.**
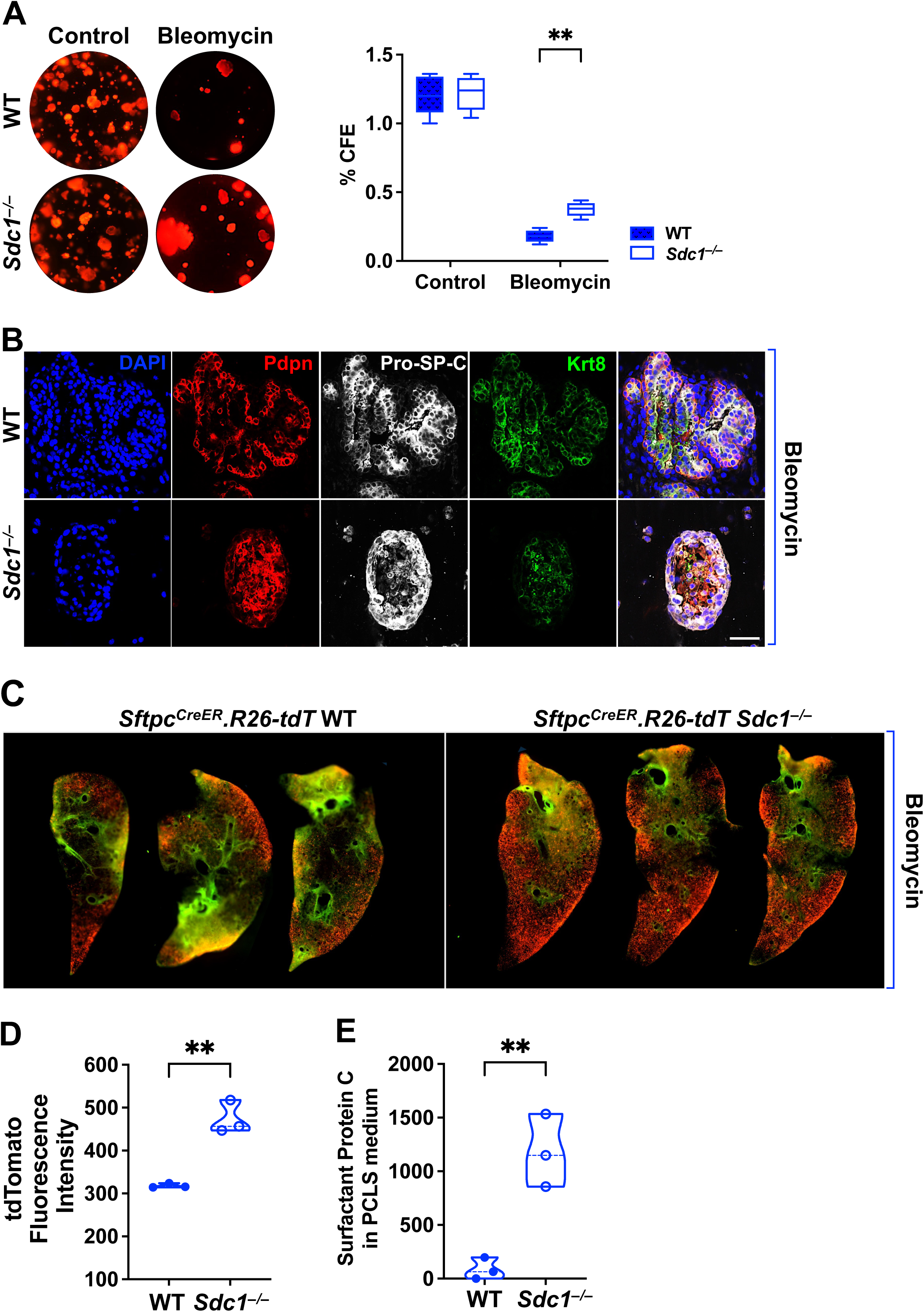
Upregulation of syndecan-1 on alveolar type 2 epithelial (AT2) hinders AT2 proliferation, differentiation, and function. **A** Representative images and colony-forming efficiency (CFE) analysis of AT2 alveolospheres derived from uninjured and day 14 bleomycin-injured WT and *Sdc1^–/–^* mice, assessed at day 21 of culture, N= 5 per group. **B** Representative immunofluorescence staining of AT2 alveolospheres derived from day 14 bleomycin-injured WT and *Sdc1^–/–^*mice, stained for AT1 cells (podoplanin [PDPN]; Alexa Fluor 594, red), AT2 cells (surfactant protein C [pro–SP-C]; Alexa Fluor 647, white), and keratin 8 (KRT8; Alexa Fluor 488, green). Scale bar=50 µm. **C** Precision-cut lung slices (PCLS) from young day 14 following bleomycin injury, comparing Sftpc.Rosa26.tdT.CreER.WT and Sftpc.*^Rosa26.tdT.CreER^*. *Sdc1^–/–^* genotype, cultured for 7 days and imaged using the Incucyte system. tdTomato (Red) fluorescence reports surfactant protein C-expressing epithelial cells, while green fluorescence reflects autofluorescence associated with mesenchymal components and lung fibrosis. N=3 per group.. **D** Quantification of tdTomato (red) fluorescence intensity in Precision-cut lung slices (PCLS) images comparing WT and *Sdc1^–/–^* mice, N=3 per group. **E** Surfactant protein C levels in precision-cut lung slice (PCLS) culture media measured by ELISA, N=3 per group. *\**p<0.05*; \*\**p<0.005*; \*\*\**p<0.0005 by Student’s t-test for two sample comparison.

### Excess syndecan-1 impairs AT2 cell function in lung tissue

Given that AT2 cells synthesize and secrete surfactant protein C (SP-C), a critical regulator of alveolar stability, and that SP-C dysfunction is linked to fibrotic lung diseases (33–36), we next assessed AT2 cell function within the lung microenvironment using precision-cut lung slices (PCLS). PCLSs were generated from uninjured and day-14 bleomycin-injured WT and *Sdc1^–/–^* Sftpc lineage-traced mice (Sftpc.CreER.R26-tdTomato; N=3 per group) and cultured in an IncuCyte S3 live-cell imaging system for 7 days. The tdTomato (Red) fluorescent intensity, representing AT2 cell abundance, was significantly higher in bleomycin-injured *Sdc1^–/–^* PCLSs than in matched WT PCLSs (**Fig. 5C–D, Supplementary Fig. S13**), whereas green autofluorescence—reflecting mesenchymal or fibrotic components—was also reduced in *Sdc1^–/–^* tissues (**Fig. 5C; Supplementary Fig. S14**). Consistent with preserved AT2 function, SP-C levels measured by ELISA in day-3 PCLS culture media were significantly higher in *Sdc1^–/–^*compared with WT slices (**Fig. 5E**). These data further support the conclusion that excess syndecan-1 impairs AT2 cell function following injury.

### Syndecan-1 induces the acetylation of p53

To identify senescence signaling pathways downstream of syndecan-1, we performed upstream regulator analysis using Ingenuity Pathway Analysis (IPA) on AT2 differentially expressed genes from IPF versus control lungs (GSE122960) (16). TP53 emerged as one of the most significantly activated transcriptional regulators in IPF AT2 cells (z-score = 2.27; adjusted P = 2.6 × 10⁻⁶²). Construction of SDC1- and TP53-centered interaction networks revealed multiple shared nodes linking the two pathways, including regulators such as CDC42 and TGFB1, implicating syndecan-1–TP53 crosstalk in senescence regulation (**Fig. 6A**, blue lines).

**Figure 6.**
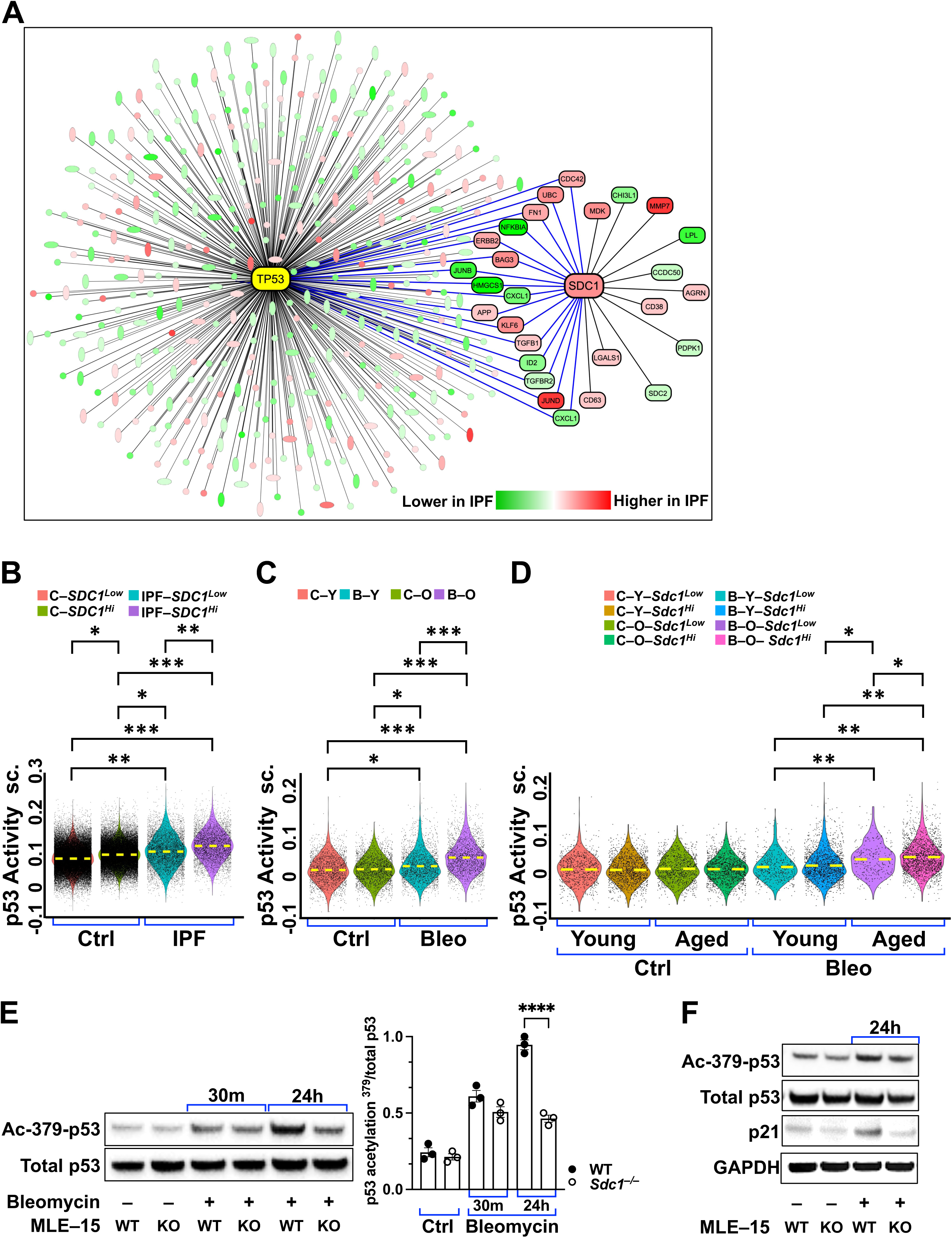
Syndecan-1 induces the acetylation of p53. **A** An interaction network analysis centered on syndecan-1 (SDC1) and TP53 was constructed from differentially expressed genes in IPF vs. control alveolar type 2 (AT2) cells of the scRNAseq dataset (**GSE122960**), highlighting shared nodes that connect both network centers (blue lines). Higher and lower gene expression in IPF is indicated by shades of red and green, respectively. **B** Violin plots showing the mean composite transcriptomic p53 activity score in alveolar type 2 (AT2) cells from control and IPF lungs in an integrative dataset of multiple published scRNA-seq cohorts (unpublished), stratified by syndecan-1 expression into high (*SDC1^Hi^*) and low (*SDC1*^low^) AT2 cell populations. **C–D** Violin plots showing the mean composite transcriptomic p53 activity score in alveolar type 2 (AT2) cells of the published dataset (**GSE186246**) of young– and aged– uninjured and bleomycin-injured WT. **(C)** Young– and aged– uninjured and bleomycin-injured WT. **(D)** Stratification of all groups according to syndecan-1 expression level as high (*Sdc1^hi^*) and low (*Sdc1^Low^*). **E–F** Representative immunoblot analysis of acetylated p53 MLE-15 cell lysates from wild-type (WT) and *Sdc1^–/–^* (KO) cells, with and without bleomycin stimulation **(E)**, and the impact of acetylation on p21 expression **(F)**. Experiments were performed in 3 independent biological replicates. *\**FDR*<*0.01 is considered significant. *\**p<0.05*; \*\**p<0.005*; \*\*\**p<0.0005 by Kruskal−Wallis, one-way ANOVA analysis for multiple comparisons, and Student’s t-test for two-sample comparisons.

We next evaluated a transcriptomic p53 activity score (**Supplementary Table 2**) in human AT2 cells using an unpublished integrative scRNA-seq cohort. AT2 cells stratified by syndecan-1 expression revealed significantly higher p53 activity scores in IPF versus control cells, with *SDC1^Hi^* AT2 cells exhibiting elevated p53 activity in both groups (**Fig. 6B**). Consistent with these findings, bleomycin-injured mouse AT2 cells exhibited increased p53 activity, with the highest scores observed in aged mice (**Fig. 6C, Supplementary Fig. S15A**). Stratification by syndecan-1 expression recapitulated the human pattern (**Fig. 6D**). Comparison of WT and *Sdc1^–/–^* AT2 cells demonstrated a trend of p53 activity attenuation in *Sdc1-*deficient cells, particularly in the aged cohort (**Supplementary Fig. S15B–C**), whereas young AT2 cells showed minimal genotype-dependent differences (**Supplementary Fig. S15B–C**). Given prior reports implicating p53 acetylation in p21 induction (37), we examined p53 acetylation at lysine 379 (K379) in WT and *Sdc1^–/–^* MLE-15 cells following bleomycin stimulation. p53^K379^ acetylation was robustly induced in WT cells but was significantly reduced in *Sdc1^–/–^* cells 24 hours after bleomycin exposure (**Fig. 6E**), without differences in total p53 abundance. Additionally, reduced p53 acetylation was associated with downregulation of p21 (**Fig. 6F**). These findings suggested that syndecan-1 promoted epithelial senescence by modulating p53 activation.

Together, these findings demonstrate that syndecan-1 promotes epithelial senescence, likely through p53-dependent signaling, impairing AT2 renewal, differentiation, and secretory function, and thereby driving maladaptive repair and fibrotic remodeling in lung disease.

## Discussion

Alveolar type 2 (AT2) epithelial cells are central to alveolar repair after lung injury, and impaired AT2 progenitor function is a defining feature of pulmonary fibrosis. In this study, we identify excess syndecan-1 as an epithelial regulator of AT2 cell–autonomous senescence that contributes to maladaptive repair and fibrotic remodeling. By integrating human transcriptomic analyses with mouse genetic models and *in vitro* epithelial systems, we find that syndecan-1 is linked to activation of p53 signaling in AT2 cells, with downstream effects on renewal capacity.

Senescent AT2 cells accumulate in IPF and experimental fibrosis and are closely associated with disease progression. In this setting, syndecan-1 is consistently upregulated in fibrotic AT2 cells across human fibrotic lung disease and mouse bleomycin injury. Functionally, excess syndecan-1 enhances senescence and disrupts key AT2 properties. These observations align with recent spatial transcriptomic studies identifying syndecan-1–enriched epithelial populations that persist in transitional, non-resolving states (38, 39). We interpret these data to suggest that syndecan-1–associated senescence constrains AT2 cells within these dysfunctional programs, thereby sustaining aberrant repair. In contrast, loss of syndecan-1 attenuates senescence, preserves features of AT2 stemness, and improves epithelial function, supporting a causal role. The reduction in surfactant protein C observed with syndecan-1 excess—whether reflecting loss of AT2 cells or impaired production—remains to be clarified. It is also plausible that syndecan-1 functions as a co-receptor for pro-senescent ligands, including insulin-like growth factor (IGF) or fibroblast growth factor (FGF) family members, although this will require direct experimental assessment.

Mechanistically, our data place syndecan-1 upstream of the p53/p21 axis. Loss of syndecan-1 reduces p53 acetylation without affecting total p53 levels, indicating regulation at the level of p53 activation rather than abundance. The mechanism underlying this effect remains to be defined. Given the cell surface localization of syndecan-1, a direct interaction with nuclear acetylation machinery is unlikely. Instead, syndecan-1 may modulate upstream signaling pathways that converge on p53. Candidates emerging from prior work and our interaction network include KLF6 (40), along with growth factor–dependent signaling pathways. Syndecan-1 binds TGF-β, EGF, and VEGF and can activate downstream cascades such as JunD and ERK/MAPK (41), which are known to intersect with p53 signaling. In this context, syndecan-1 may act as a signaling node that amplifies extracellular stress cues, enhancing p53 acetylation and reinforcing a senescence program in epithelial cells.

The role of syndecan-1 in senescence is likely context dependent, with prior studies reporting divergent effects in fibroblasts (42, 43). Our findings instead support a pro-senescent role for excess intact syndecan-1 within the epithelium in fibrotic lung disease. Given the central contribution of AT2 senescence to fibrosis pathogenesis, modulation of syndecan-1 may represent a strategy to restore epithelial repair capacity. Future studies using AT2-specific genetic approaches and ex vivo human lung models will be important to determine whether targeting syndecan-1 is therapeutically feasible.

## Methods

### Cell lines and 3D mouse AT2 alveolosphere cultures

The following cell lines: mouse lung epithelial cells (MLE-15; Applied Biological Materials, British Columbia, Canada), human bronchial epithelial cells (BEAS-2B; ATCC, Manassas, VA), and human alveolar epithelial carcinoma cells (A549; ATCC) were maintained and propagated according to established protocols. MLE-15 cells were maintained in DMEM/F-12 (1:1; Corning, Salt Lake City, UT) supplemented with 2% fetal bovine serum (FBS; HyClone, Logan, UT), 1% penicillin–streptomycin (Corning), 10 nM β-estradiol (MilliporeSigma, Burlington, MA), 10 nM hydrocortisone (MilliporeSigma), and insulin–transferrin–selenium (ITS; Gibco, Thermo Fisher Scientific, Waltham, MA). BEAS-2B cells were cultured in BEGM basal medium supplemented with the Bronchial Epithelial Cell Growth Medium BulletKit (Lonza, Morristown, NJ). A549 cells were maintained in DMEM supplemented with 10% FBS and 1% penicillin–streptomycin. All cell lines were cultured at 37°C in a humidified incubator with 5% CO₂.

CRISPR/Cas9-mediated deletion of syndecan-1 (SDC1 knockout; KO) was performed in MLE-15 and A549 cells by Synthego Corporation (Redwood City, CA, USA). Lentiviral transduction was used to overexpress mouse syndecan-1 (OE) in MLE-15 cells and to knock down syndecan-1 (SDC1 KD) in BEAS-2B cells using shRNA targeting SDC1, with re-expression of mouse syndecan-1 performed under knockdown conditions as indicated. Empty vector controls and non-targeting guide RNA constructs were used for lentiviral and CRISPR experiments, respectively. Syndecan-1 expression levels in all cell lines were confirmed by flow cytometry.

Three-dimensional (3D) AT2 alveolosphere cultures were generated and maintained according to established protocols (44). Briefly, AT2 cells were isolated from SftpcCreER-R26-tdTomato WT and *Sdc1^–/–^* mouse lungs using enzymatic lung digestion as previously described (44, 45). Single-cell suspensions were stained with lineage-specific surface markers, and AT2 cells were purified by fluorescence-activated cell sorting as DAPI⁻CD31⁻CD45⁻EpCAM⁺tdTomato⁺ cells. Freshly sorted AT2 cells were mixed with mouse lung fibroblasts (MLg; ATCC) and growth factor–reduced Matrigel (Corning Matrigel Basement Membrane Matrix, GFR) at a 1:1 ratio and plated onto 0.4-μm transwell inserts. Alveolospheres were cultured with AT2 growth medium (DMEM/F-12 supplemented with 10% active FBS, ITS, 1% Antibiotic–Antimycotic, and the TGF-β receptor inhibitor SB-431542; MilliporeSigma) in the lower compartment of 24-well transwell plates. Cultures were maintained at 37°C with 5% CO₂, and colony-forming efficiency (CFE) was quantified by serial imaging using a Nikon Eclipse Ti-E microscope (Nikon Instruments Inc., Melville, NY, USA).

### Animals

All animal experiments were approved by the Institutional Animal Care and Use Committee (IACUC) at Cedars-Sinai Medical Center (protocol #00009018) and conducted in accordance with NIH guidelines. C57BL/6 wild-type (WT) and global syndecan-1–deficient (*Sdc1^–/–^*) mice were maintained under specific pathogen-free conditions. Young mice were 8–12 weeks of age, and aged mice were 18–24 months of age. All mice were derived from in-house breeding colonies. Male mice were used for all experiments according to our preliminaty data indicated significant lung fibrosis in male mice. Mice were randomly assigned to experimental groups.

### Bleomycin-induced lung injury and fibrosis model

Pulmonary fibrosis was induced using a single intratracheal administration of bleomycin sulfate (0.25–0.75 U/kg; APP Pharmaceuticals, Schaumburg, IL, USA) following isoflurane anesthesia. Sterile Dulbecco’s phosphate-buffered saline (DPBS) was used as a vehicle control. Bleomycin dosing was selected based on age, genotype, and experimental endpoint and is consistent with established protocols for inducing reproducible lung injury and fibrosis while minimizing mortality in young and aged mice. Animals were monitored daily for clinical signs of distress, including changes in activity, grooming, and body weight. Bronchoalveolar lavage fluid (BALF) collection and lung tissue harvesting were performed at days 14, 21, or 28 after bleomycin administration, depending on the experimental design. Lung tissues were processed for downstream analyses, including enzymatic digestion for single-cell suspension preparation, protein extraction, histological assessment, and hydroxyproline quantification.

### Precision–Cut Lung Slices (PCLS)

Precision-cut lung slices (PCLS) were generated from uninjured (control) and bleomycin-injured Sftpc.CreER-R26-tdTomato WT and *Sdc1^–/–^*mice according to a previously published protocol (46). Lung tissue sectioning was performed using a Precisionary Compresstome (VF-510-0Z; Precisionary Instruments, Ashland, MA). PCLS were cultured in phenol red–free DMEM (Thermo Fisher Scientific) supplemented with 0.1% fetal bovine serum (FBS) and 1% penicillin–streptomycin. Cultures were maintained in an IncuCyte live-cell imaging humidified incubator at 37°C with 5% CO₂ and subjected to serial live-cell imaging for up to 7 days. Immunofluorescence staining of PCLS was performed as previously described(46).

### Protein assays, histology, and immunostaining

Protein concentrations from cell lysates, tissue homogenates, and culture supernatants were quantified using the bicinchoninic acid (BCA) protein assay (Pierce, Thermo Fisher Scientific) according to the manufacturer’s instructions. Western blotting, enzyme-linked immunosorbent assays (ELISA), and hydroxyproline assays were performed as previously described (30).

Formalin-fixed, paraffin-embedded human and mouse lung tissues were sectioned at 4–10 μm thickness and processed for histological analyses using standard protocols. Briefly, sections were air-dried to promote tissue adhesion and baked at 60°C for 2 hours prior to staining. After deparaffinization and rehydration, antigen retrieval was performed by steaming slides in 1× citrate buffer (pH 6.0) for 20 minutes. Sections were permeabilized with 0.2% Triton X-100 in PBS for 10 minutes and blocked with 5% bovine serum albumin (BSA) in PBS for 45 minutes. Slides were incubated overnight at 4°C with primary antibodies, followed by incubation with appropriate secondary antibodies the next day. Hematoxylin and eosin (H&E), Masson’s trichrome, Picrosirius red, and immunofluorescence staining were performed according to established laboratory protocols (30). All antibody clones and other details, including sources and catalog numbers, are listed in **Supplementary Table 3.** Images were acquired using a Nikon Eclipse Ti2 microscope (Nikon Instruments Inc., Melville, NY, USA) or an Axioscan 7 slide scanner (Carl Zeiss Microscopy, Oberkochen, Germany) at 20x magnification.

### Image analysis and quantifications

Whole–tissue section image analysis was performed using Nikon NIS-Elements software with the General Analysis 3 (GA3) module and QuPath (version 0.6.0(47). For analysis in Nikon GA3, lookup table (LUT) settings were optimized to minimize background fluorescence and were applied uniformly across all images prior to quantification. Images were then processed using the Rolling Ball background subtraction and Auto Contrast functions to correct for uneven illumination and enhance object detection. Cells were identified in each fluorescence channel using Bright Spots detection and filtered based on cell size, local contrast, and signal intensity thresholds. Speckles were removed using the Clean function, and cellular objects were further segmented and contoured using the Grow Regions, Grow Objects, and Make Cell functions, with regions of interest (ROIs) selectively retained based on overlap with DAPI-positive nuclei. Total cell counts by fluorescent marker, along with per-cell mean and maximum fluorescence intensities, were exported for downstream analysis. QuPath was also used for whole-slide image analysis and cell segmentation. Cell classifiers were trained on representative regions and applied across entire tissue sections to determine marker-specific cell counts.

### Single–cell RNA sequencing (scRNAseq)

The young control and bleomycin-injured WT and *Sdc1^–/–^* cohort details were published (30). For the aged mouse cohort, freshly harvested bleomycin-injured WT and *Sdc1^–/–^* mouse lungs were enzymatically digested with elastase (Worthington Biochemical Corporation, Lakewood, NJ) and Liberase (RocheTM, Millipore-Sigma), and then mechanically disrupted to generate a single–cell suspension, as previously published (16). Viable lung cell suspension (DAPI^–^) was flow-sorted for lung immune cells (CD45^+^), endothelial cells (CD31^+^), epithelial cells (CD326^+^), and stromal cells (CD45^–^CD31^–^CD326^–^). Equal proportions of all four cell types were pooled and subjected to single-cell capture using Evercode^TM^ WT mini kit v2 for barcoding and library preparation (Parse Bioscience, Seattle, WA) following the manufacturer’s protocol. Sequencing was performed using an Illumina Novaseq SP at 200 cycles, with 800 million reads (Illumina, San Diego, CA), at the Cedars-Sinai Medical Center, Applied Genomic, Computational, and Translational (AGCT) Core. ParseBiosciences–Pipeline 1.1.2 was used with the default settings for demultiplexing and aligning reads to the mouse GRCm39 reference genome.

### RNAseq data analysis

Single-cell analysis R package Seurat v5.0 was used for data analysis (48). To ensure quality and filter out low-quality cells, only cells expressing more than 200 genes (defined as genes detected in at least 3 cells) and fewer than 10% mitochondrial genes were selected. To minimize doublet contamination in each dataset, the Doubletfinder pipeline was used to remove potential doublets, with the doublet ratio determined using a fit model generated from the suggested “multiplet rate”/“number of cells recovered” ratio, as in the 10X Genomics user manual (49). The batch correction package Harmony was used for data integration (50). The batch correction was performed using Principal Component Analysis (PCA) on the 3000 most variable genes, and the first 30 independent components were used for downstream, unbiased clustering with a resolution of 0.6. The Uniform Manifold Approximation and Projection (UMAP) method was used for unsupervised clustering visualization. Cell cluster identities were determined using known gene markers of major immune cell types (51). Differentially expressed genes between different clusters and groups were calculated using Mode-based Analysis of Single-cell transcriptomics (MAST) (52). Differentially expressed genes with adjusted p-value <0.05 were used for downstream analysis. Functional enrichment analysis using KEGG pathways was performed using WebGestalt, as described (53).

### Upstream regulator and network analysis

Ingenuity Pathway Analysis (IPA, Qiagen, CA) was used to identify upstream regulators based on scRNAseq differentially expressed (adjusted *P*-value *<* 0.05) gene patterns of AT2 cells in IPF and control samples. IPA’s knowledge base was used to generate interaction networks using syndecan-1 and TP53 as seeds.

### Spatial transcriptomic sequencing and analysis

The spatial transcriptomic dataset can be accessed through the Gene Expression Omnibus (GEO) under accession number GSE292956. Samples were prepared and processed using the Visium 10x protocol (10X Genomics, Pleasanton, CA). Tissue samples were prepared and processed using the Visium Spatial Gene Expression protocol (10x Genomics, Pleasanton, CA). Primary data processing was performed using the 10x Genomics Space Ranger pipeline. Downstream analyses were conducted using Partek Flow software (Illumina, San Diego, CA).

Spatial transcriptomic data were analyzed using Partek Flow software (Illumina, San Diego, CA), which was used to process Space Ranger output and perform downstream normalization, dimensionality reduction, clustering, and differential expression analyses. Data were normalized and scaled using term frequency–inverse document frequency (TF-IDF). Dimensionality reduction was performed using principal component analysis (PCA), t-distributed stochastic neighbor embedding (t-SNE), and uniform manifold approximation and projection (UMAP). Batch effects were addressed using Seurat v3 integration. Cell clustering was performed using k-means, graph-based, and hierarchical clustering approaches, followed by biomarker-based computational analysis for cluster annotation. Differential gene expression analyses were conducted using DESeq2 and nonparametric statistical tests, including Kruskal–Wallis followed by Dunn’s multiple-comparison tests, as appropriate. Student’s t tests were used for pairwise comparisons of mean values where applicable.

### Statistical analysis

All data are presented as mean ± SEM unless otherwise indicated. Statistical analyses were performed using GraphPad Prism (version 9 or later) and R, as appropriate. Comparisons between two groups were performed using unpaired two-tailed Student’s t tests or nonparametric equivalents when data did not meet normality assumptions. Comparisons among multiple groups were analyzed using one- or two-way ANOVA with appropriate post hoc multiple-comparison tests, or Kruskal–Wallis tests followed by Dunn’s correction, as indicated. For transcriptomic analyses, differential gene expression was assessed using DESeq2, and statistical significance was defined using a false discovery rate (FDR) threshold of < 0.01 unless otherwise stated. No data were excluded unless technically compromised. Sample sizes, biological replicates, and statistical tests used for each experiment are specified in the corresponding figure legends.

### Study Approvals

Human lung tissue was collected, processed, and stored at Cedars-Sinai Medical Center (CSMC; Los Angeles, CA) in accordance with guidelines and with approval from the CSMC Institutional Review Board (IRB# Pro00035409). Animal experiments were conducted in accordance with and approved by the Institutional Animal Care and Use Committee (IACUC) at CSMC (IACUC#000009018).

## Author Contributions

CY, ME, XL, TP, DJ, PWN, CH, BRS, WCP, and PC contributed to the conceptual design of all experiments. TP, CY, XL, YW, ZO, SC, KM, AS, and EB performed experiments. CY, ME, XL, CLM, SAG, and TP analyzed data. MZ, VK, XZ, LLP, YW, SJ, AC, FC, JL, DJ, CME, DJ, CH, NB, and BRS provided material and technical assistance. TP, CY, ME, XL, PWN, WCP, SAG, and PC were involved in writing the manuscript.

## Acknowledgments

The work is supported by the NIH grants R01HL174973 (CY), HL120947 (PC), HL103868 (PC), HL137076 (PC), AI137111 (SAG), K08 HL141590 (TP), 1K01AG090692 (LPL), T32HL170963-01A1 (AS), and California Institute for Regenerative Medicine (CIRM EDUC4-12751) (AS).

## Table Legends

**Table 1. Human SenMayo Score Gene list.**

**Table 2. Mouse SenMayo Score Gene list.**

**Table 3. Human p53 Activity Score Gene list.**

**Table 4. Mouse p53 Activity Score Gene list.**

**Table 5. Antibodies and Reagents**

## Supplementary Figures

**Figure S1.**
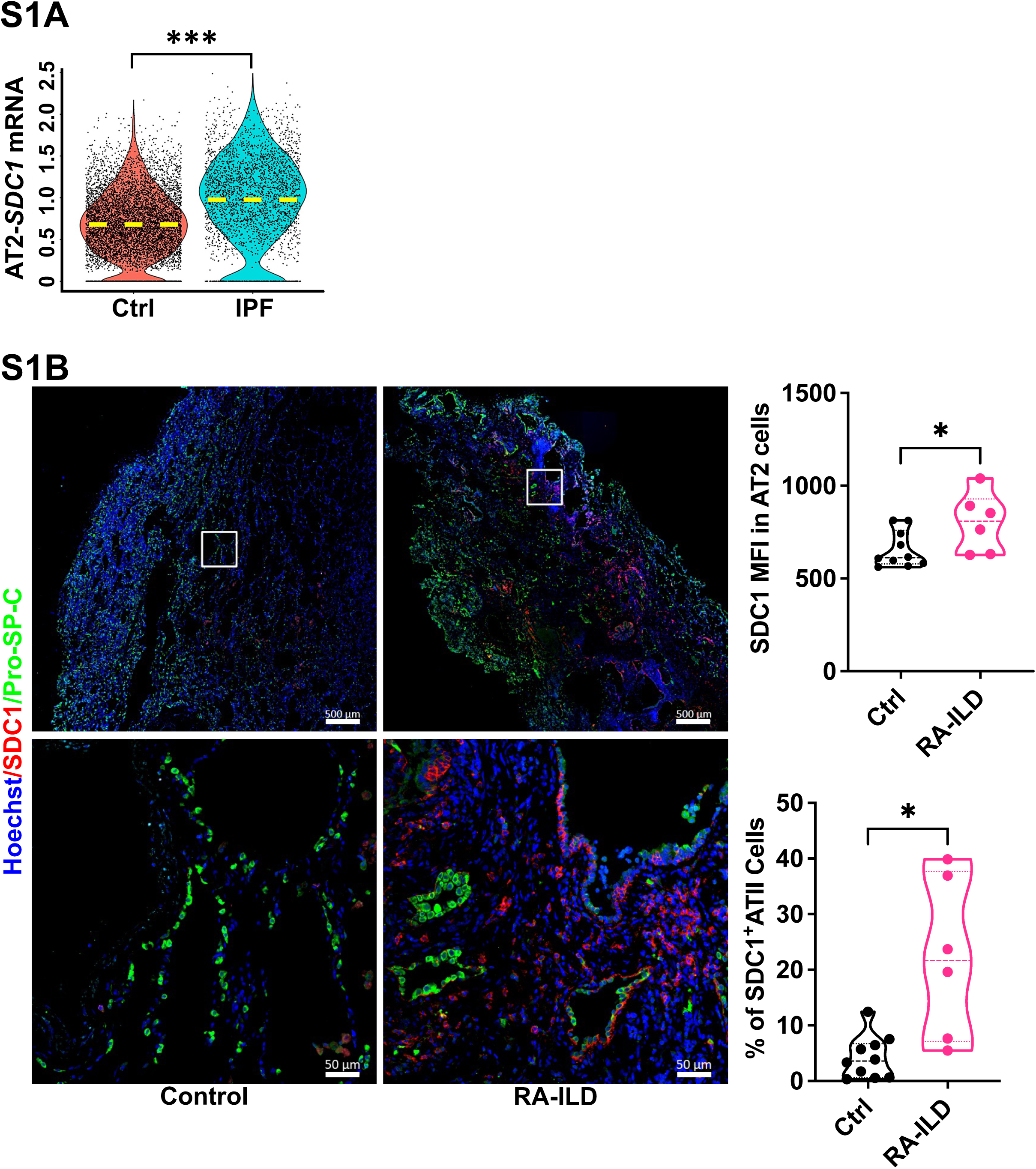
S1A Violin plot depicted syndecan-1 (*SDC1*) expression on AT2 cells of controls (N=80) vs. IPF (N=85) (unpublished cohort). **S1B** Immunofluorescence staining of syndecan-1 (SDC1, Alexa Fluor 568-red) and prosurfactant protein C (Pro-SP-C, Alexa Fluor 750-green) of control (N=10) and rheumatoid arthritis-related interstitial lung disease (RA-ILD) (N=6) human explanted lungs. Images were representative staining of an RA-ILD and control lung. Image quantification using QuPath analysis software for mean fluorescent intensity (MFI) of syndecan-1 on AT2 cells (Pro-SP-C (+)) (top) and median with interquartile range of dual syndecan-1 and Pro-SP-C (+) cells frequencies (bottom) within the whole tissue section. Scale bar=50 μm. *p<0.05; **p<0.005; ***p<0.0005 by Kruskal−Wallis, one-way ANOVA analysis.

**Figure S2.**
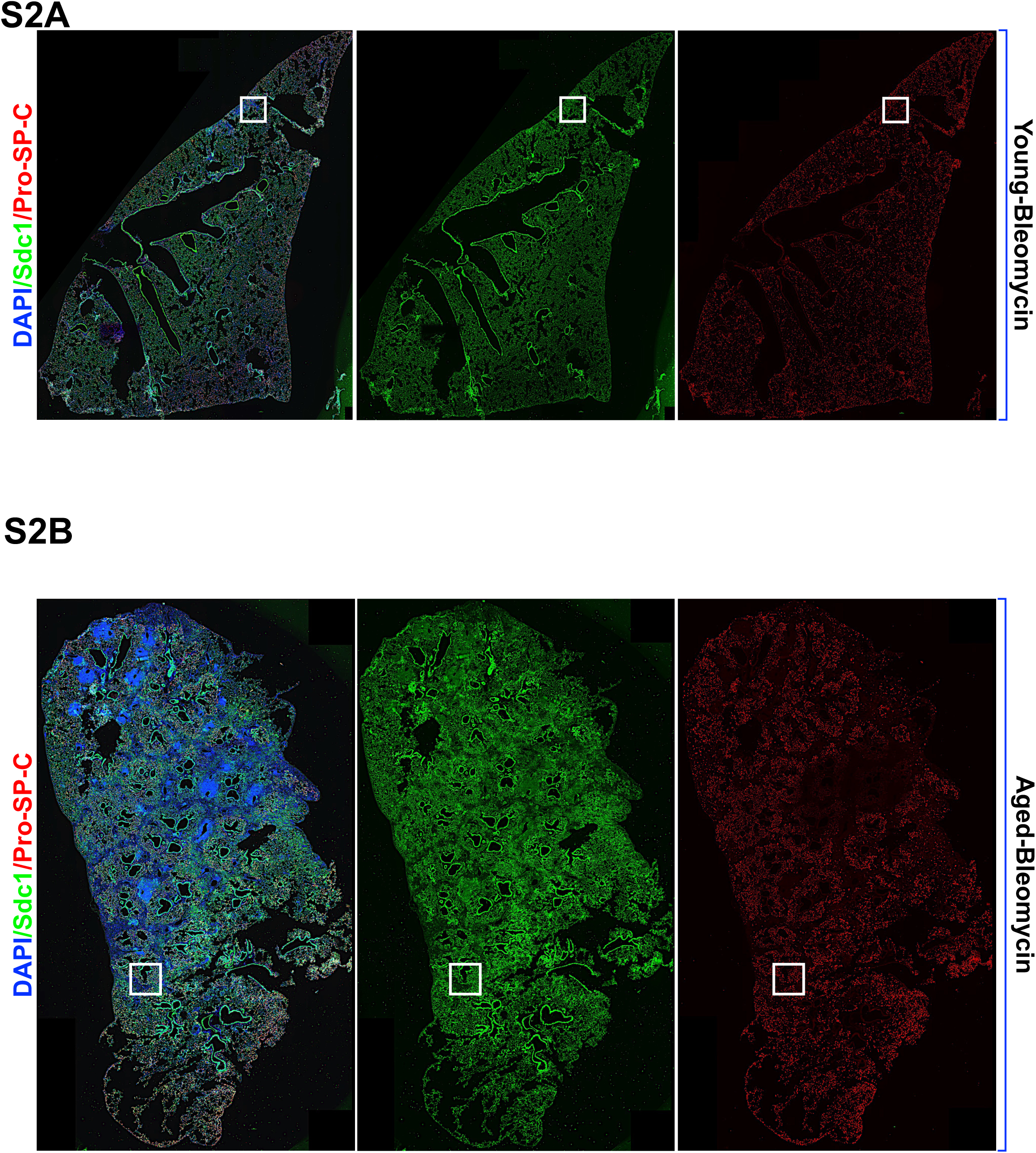
S2A–B Representative immunofluorescence whole lung section images of young **(S2A)** and aged **(S2B)** WT bleomycin-injured mouse lungs from **Fig. 2B** in individual channel for syndecan-1 (Sdc1, Alexa Fluor 488-green), and AT2 cells (Pro-SP-C, Alexa Fluor 555-red).

**Figure S3.**
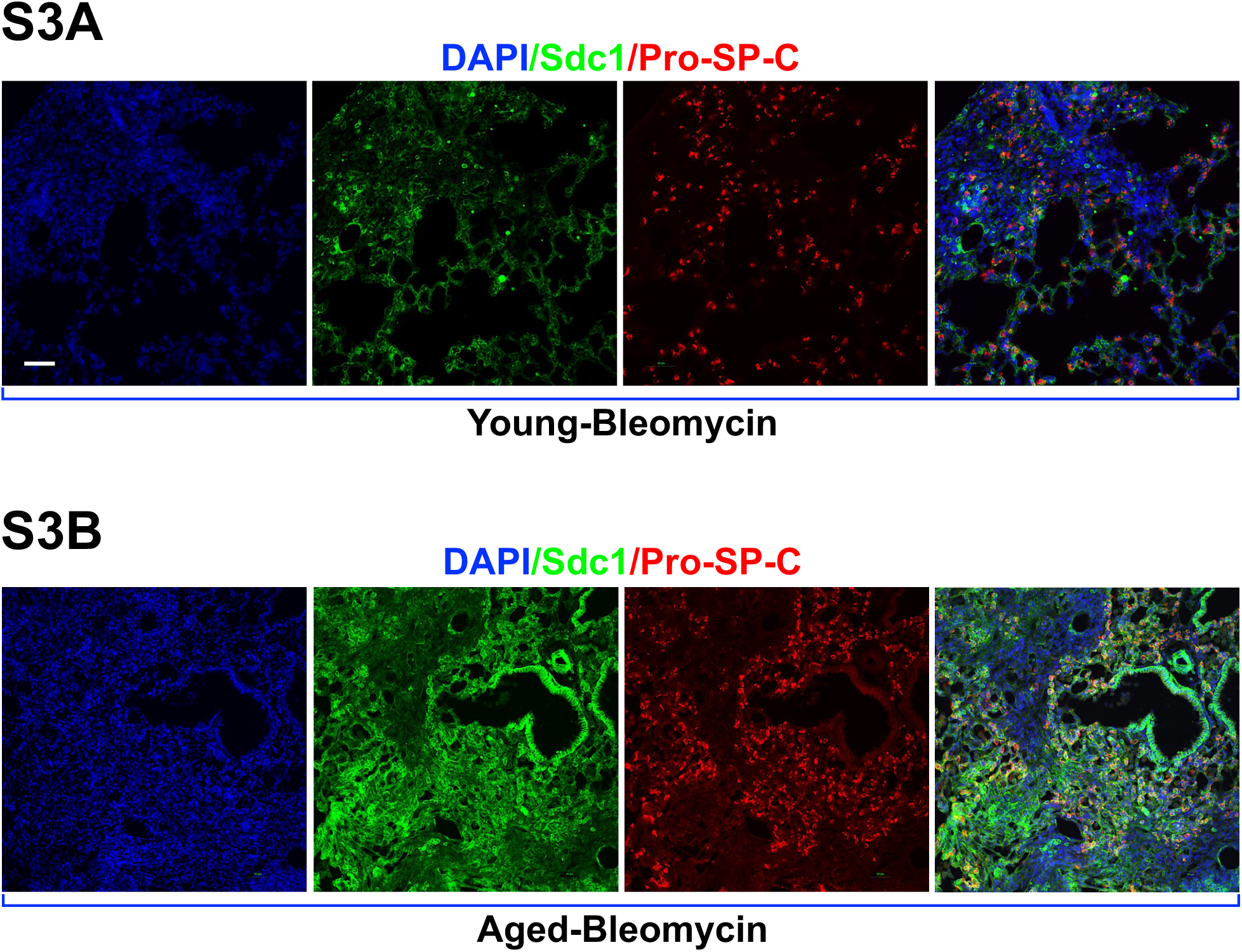
**S3A–B** Representative immunofluorescence images of young **(S3A)** and aged **(S3B)** WT bleomycin-injured mouse lungs of **Fig. 2B** in individual channel for syndecan-1 (Sdc1, Alexa Fluor 488-green), and AT2 cells (Pro-SP-C, Alexa Fluor 555-red). Scale=50 µm.

**Figure S4.**
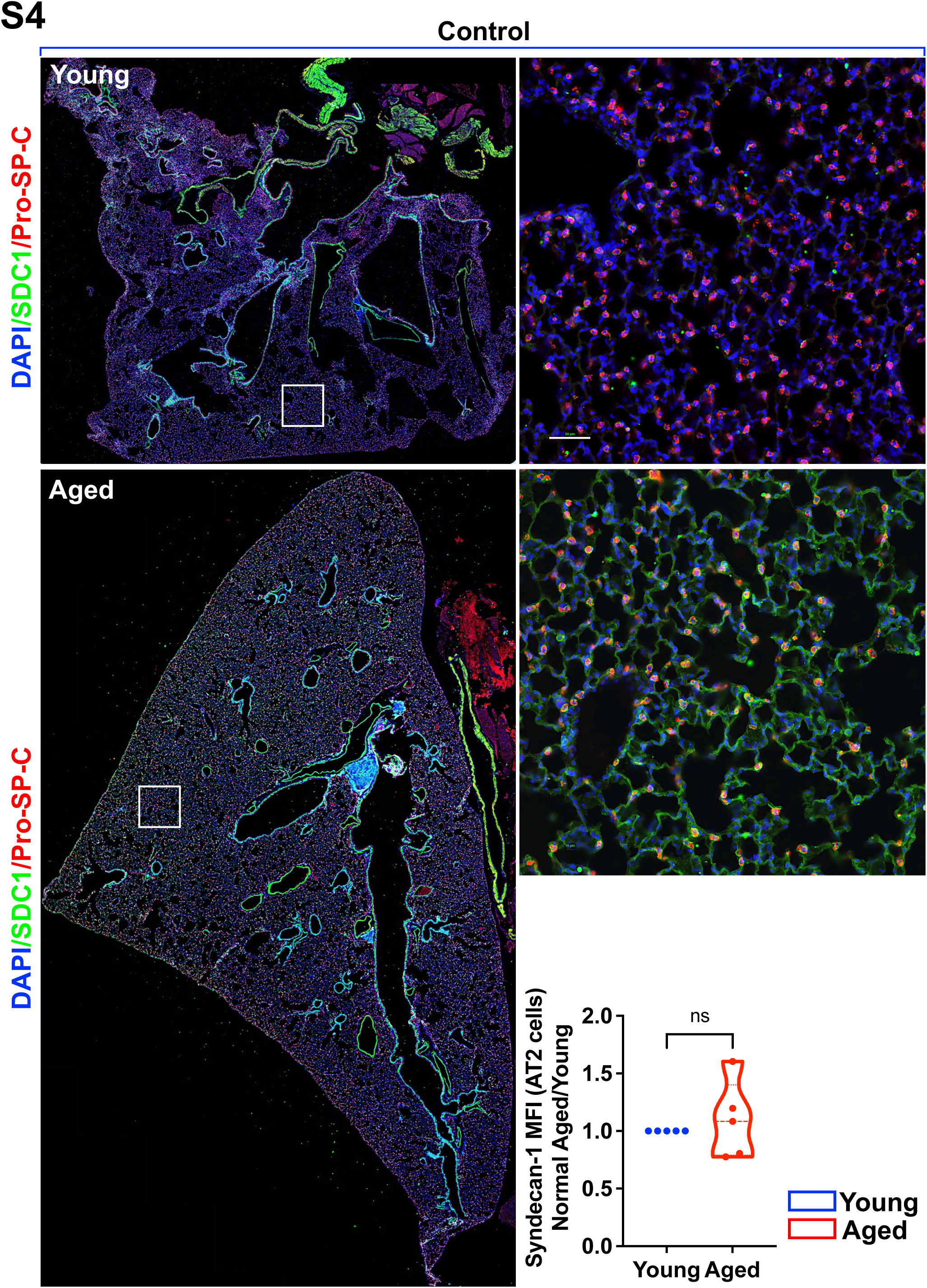
S4 Representative immunofluorescence whole lung section and selective high magnification images of young and aged WT uninjured (control) mouse lungs for syndecan-1 (Sdc1, Alexa Fluor 488-green) and AT2 cells (Pro-SP-C, Alexa Fluor 555-red. Scale=50 µm. Image quantification using Nikon GA3 for syndecan-1 mean fluorescence intensity (MFI) in pro–SP-C⁺ AT2 cells across whole-lung sections from uninjured young and aged mice (N=3 each). *p<0.05; **p<0.005; ***p<0.0005 by Student’s t-test for two sample comparison.

**Figure S5.**
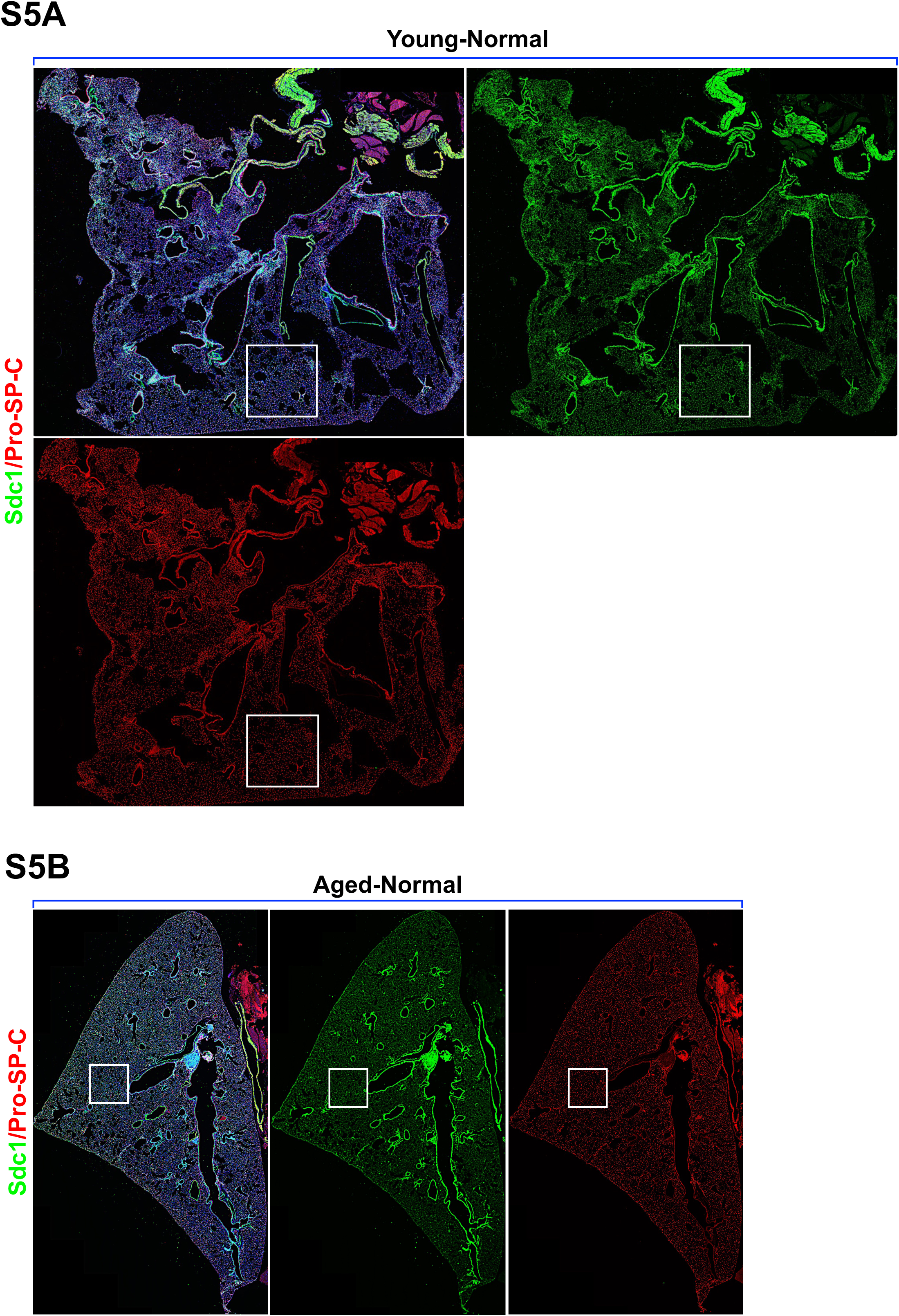
S5A–B Representative immunofluorescence whole lung section images of young **(S5A)** and aged **(S5B)** WT bleomycin-injured mouse lungs of **Supplementary Fig. 4A** in an individual channel for syndecan-1 (Sdc1, Alexa Fluor 488-green) and AT2 cells (Pro- SP-C, Alexa Fluor 555-red).

**Figure S6.**
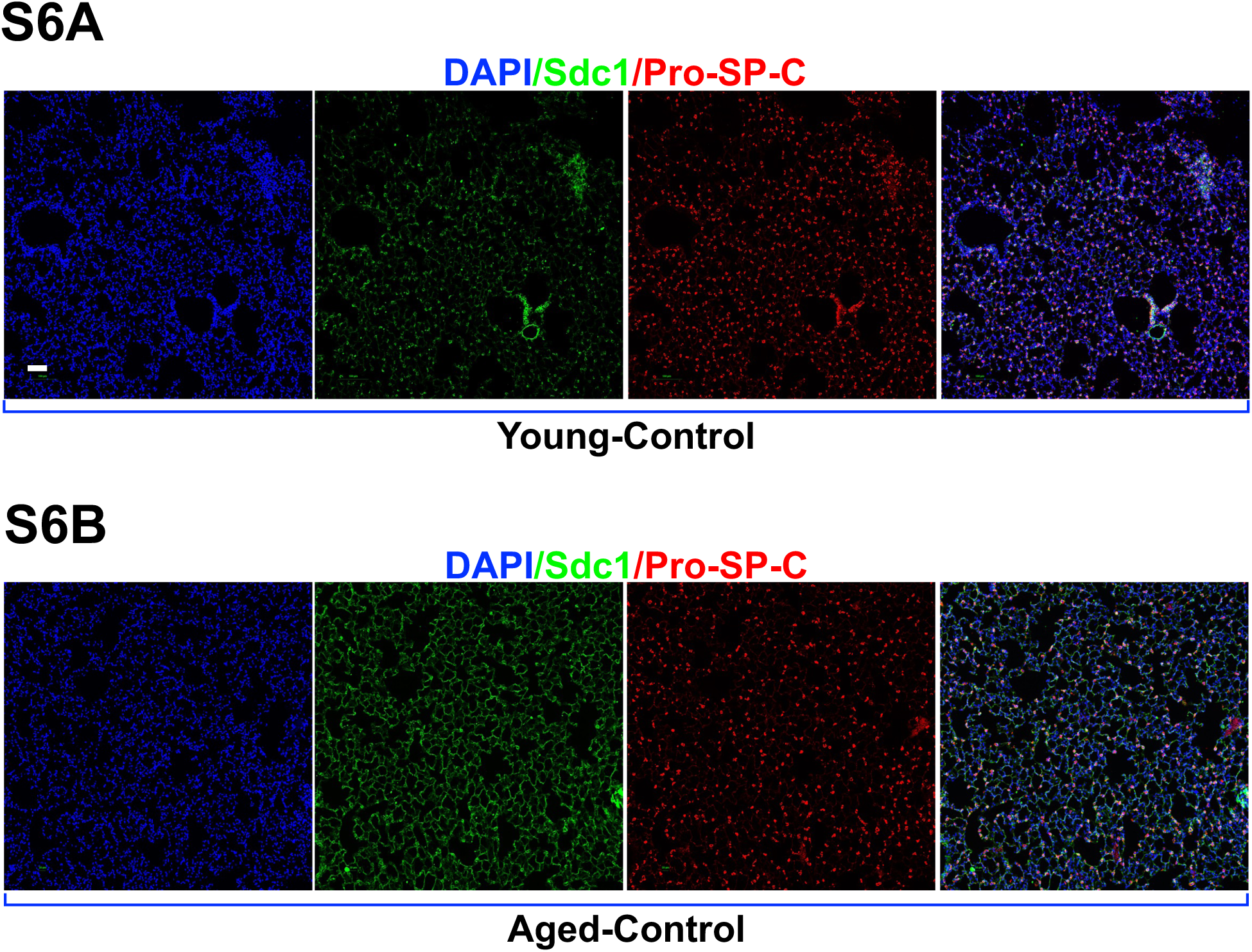
S6 Representative immunofluorescence images of young **(S6A)** and aged **(S6B)** WT bleomycin-injured mouse lungs of **Supplementary Fig. 5A** in an individual channel for syndecan-1 (Sdc1, Alexa Fluor 488-green) and AT2 cells (Pro-SP-C, Alexa Fluor 555-red) of zoomed areas (white square region). Scale=50 µm.

**Figure S7.**
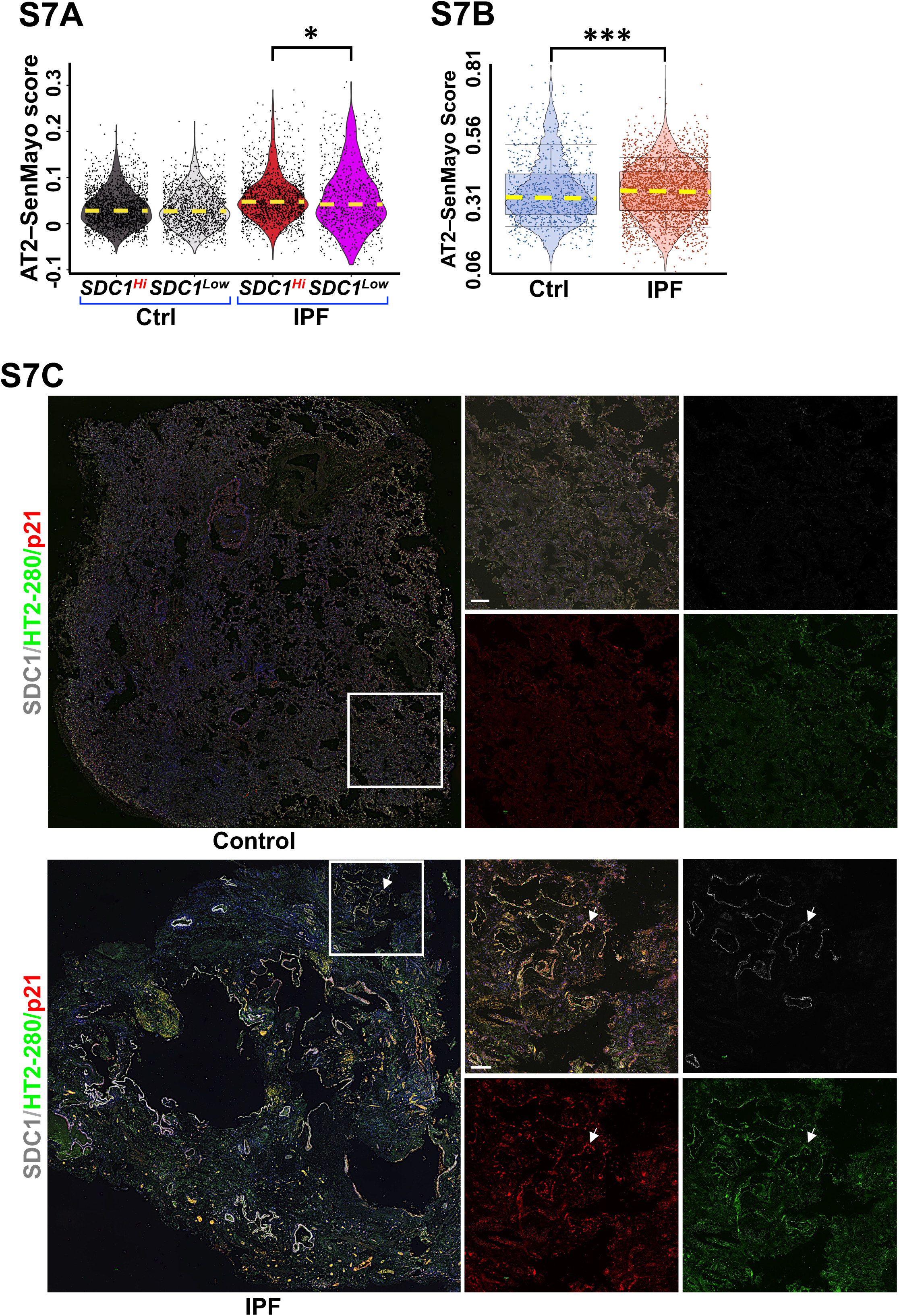
S7A Violin plot showing SenMayo senescence score of AT2 cells from controls and IPF (**GSE122960**) stratified by syndecan-1 expression into high (SDC1^Hi^) and low (SDC1^Low^) AT2 cell populations. **S7B** Violin plot depicted SenMayo senescence score of control (N=3) vs IPF (N=2) from of the spatial transcriptomic dataset (**GSE292956**). **S7C** Representative immunofluorescence images of control (N=3) and IPF (N=3) lung sections stained for AT2 cells syndecan-1 (SDC1; Alexa Fluor 647, white), AT2 cells (HT2-280; Alexa Fluor 488, green) and CDKN1A (p21; Alexa Fluor 546, red) Quantification of % p21^+^ on SDC1^+^ AT2 cells were performed in the lung sections of 3 controls and 3 IPF lungs. Scale bar=100 μm. *p<0.05; **p<0.005; ***p<0.0005 by Kruskal−Wallis, one-way ANOVA analysis.

**Figure S8.**
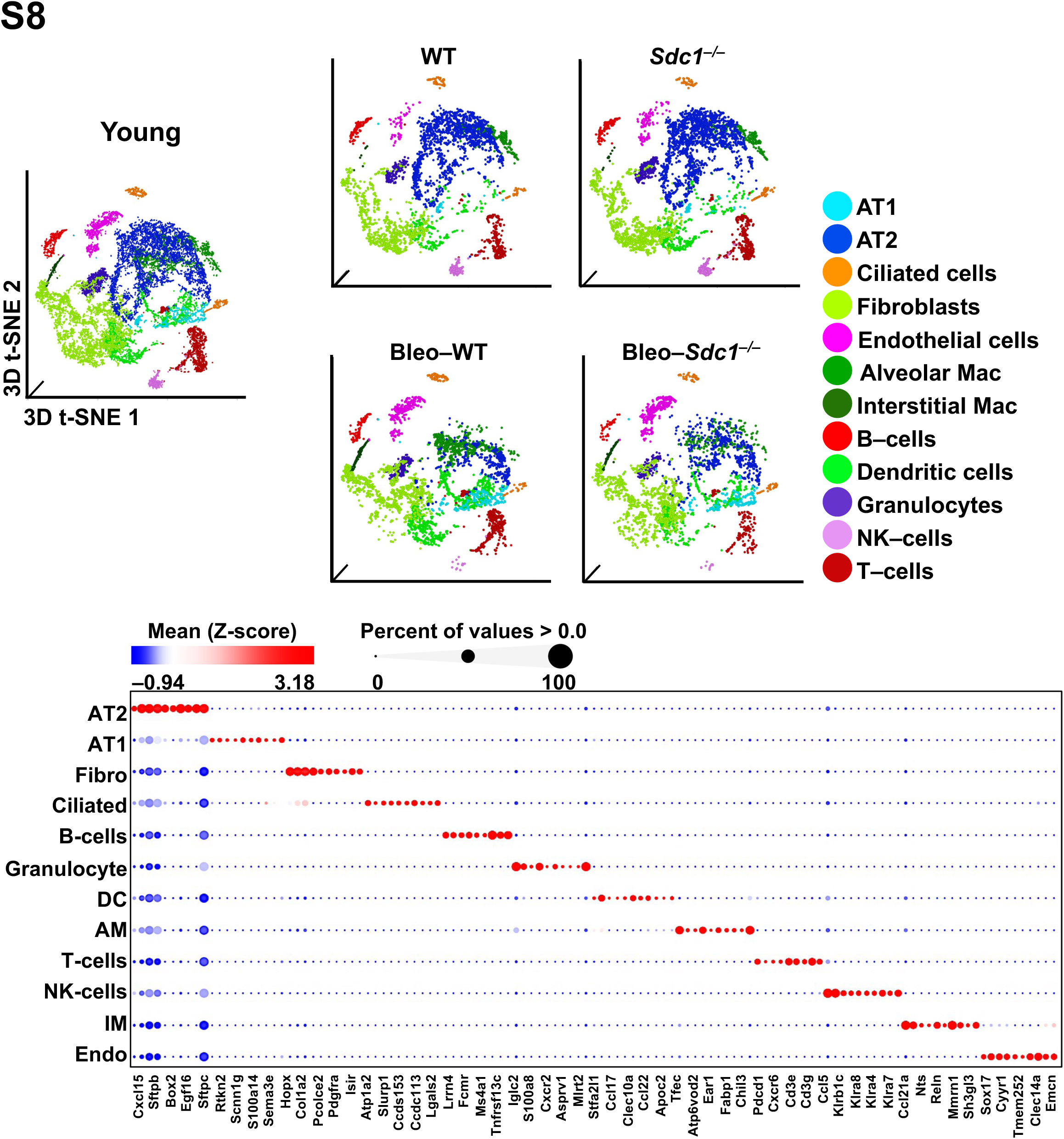
S8 t-SNE and bubble plots illustrating cellular cluster annotation in lung single cells from uninjured and bleomycin-injured young–WT and –*Sdc1^–/–^* murine dataset (**GSE134948**) (N=3 per group).

**Figure S9.**
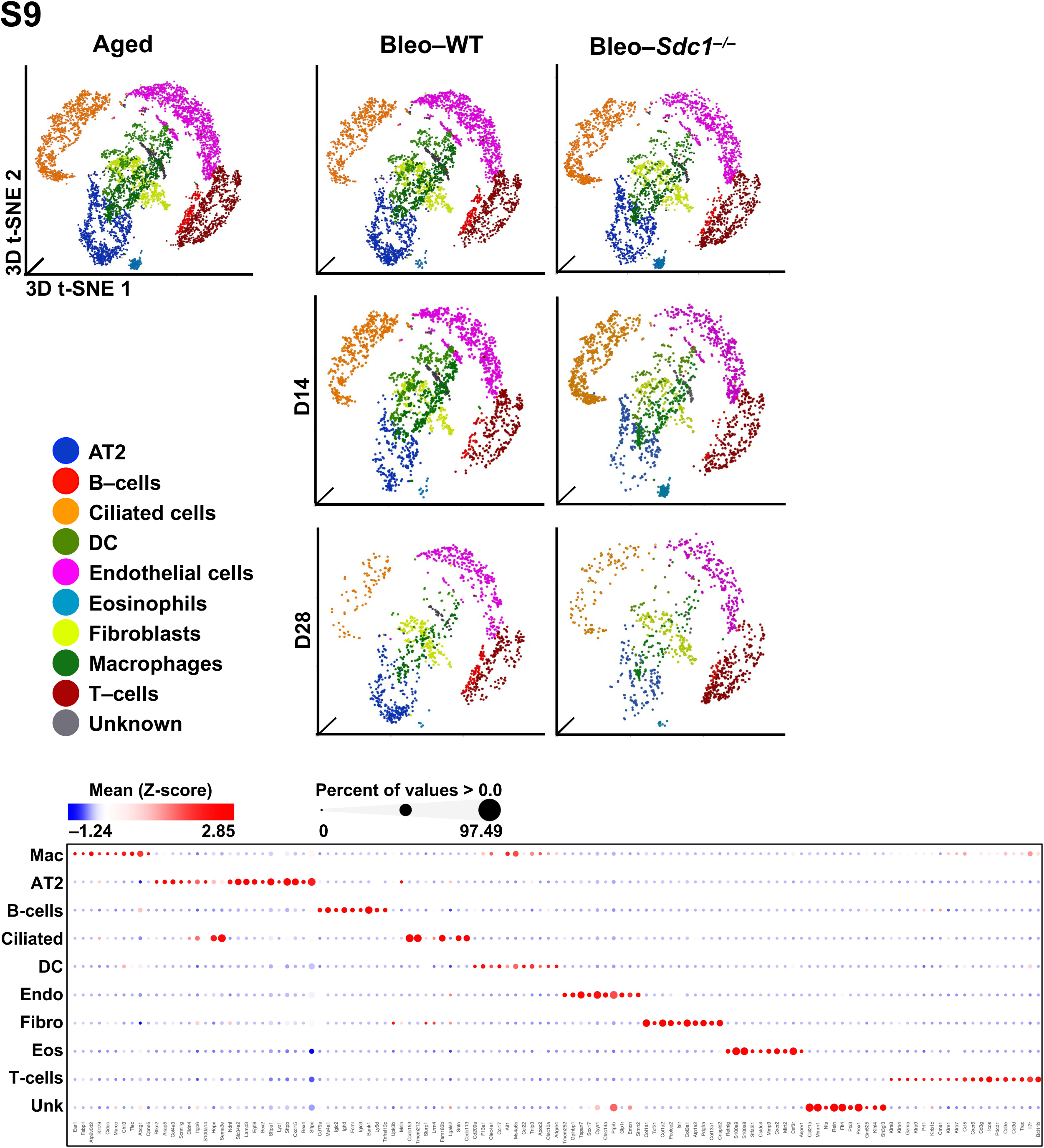
S9 t-SNE and bubble plots illustrating cellular cluster annotation of lung single cells from uninjured and bleomycin-injured aged–WT and –*Sdc1^–/–^* murine dataset (**GSE292961**) (N=3 in each group).

**Figure S10.**
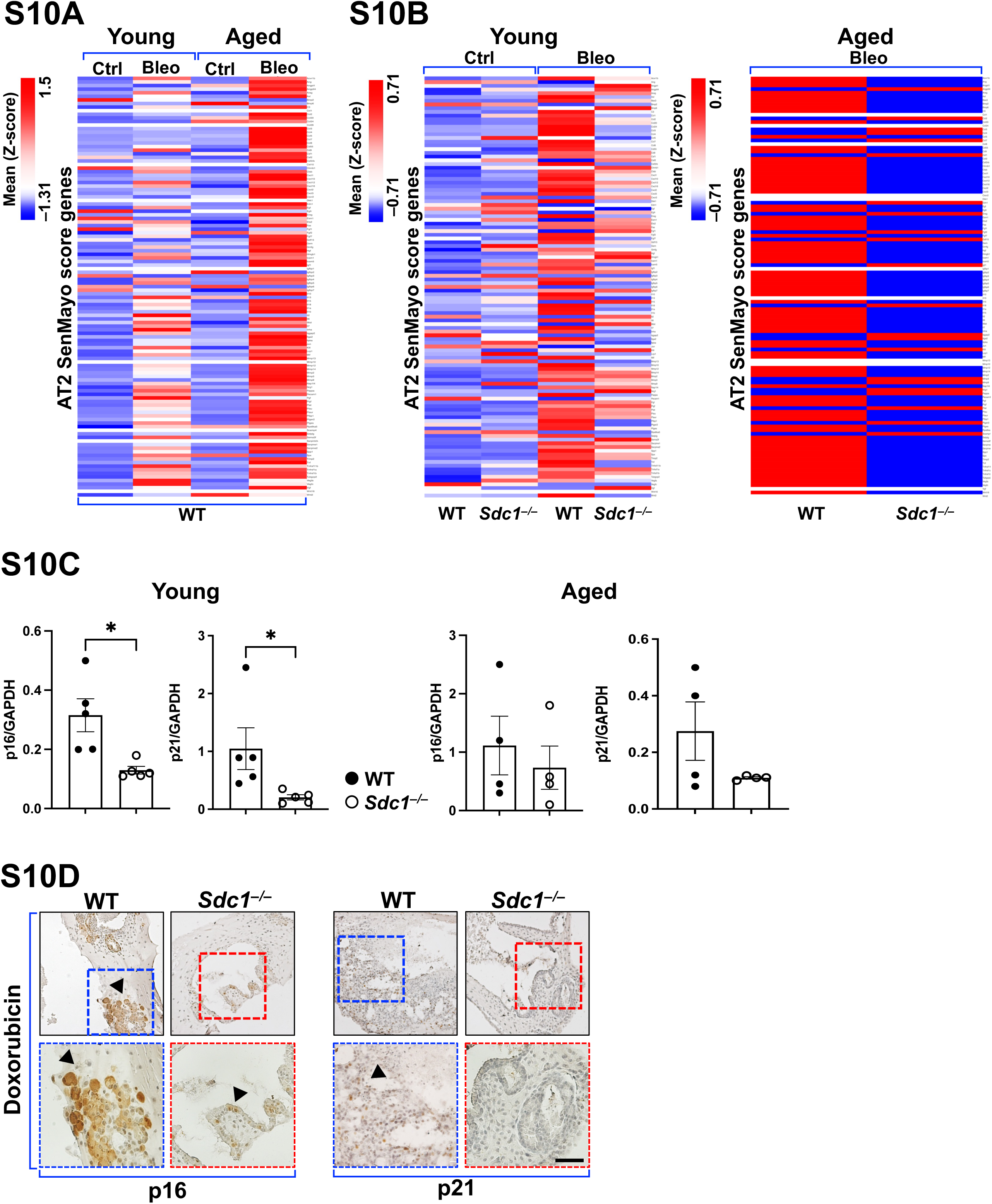
S10A–B Heatmap illustrated gene expression profiles contributing to the SenMayo senescence score of AT2 cells. **(A)** AT2 cells from young and aged WT mice under uninjured and bleomycin-injured conditions (**GSE186246**). (**B)** AT2 cells from young uninjured and bleomycin-injured WT and *Sdc1^–/–^*mice (**GSE134948**) and aged bleomycin injured WT and *Sdc1^–/–^* mice (**GSE292961**). **S10C** Densitometric quantification of p16 and p21 immunoblot signals in whole-lung homogenates from young and aged mice following bleomycin-induced lung fibrosis. **S10D** Representative immunostaining for senescence markers p16 and p21 in AT2 alveolospheres derived from young WT and *Sdc1^–/–^* mice following doxorubicin stimulation. Brown signal indicates p16 or p21 expression. Scale bar=1000 μm. *p<0.05; **p<0.005; ***p<0.0005 by Kruskal−Wallis, one-way ANOVA analysis.

**Figure S11.**
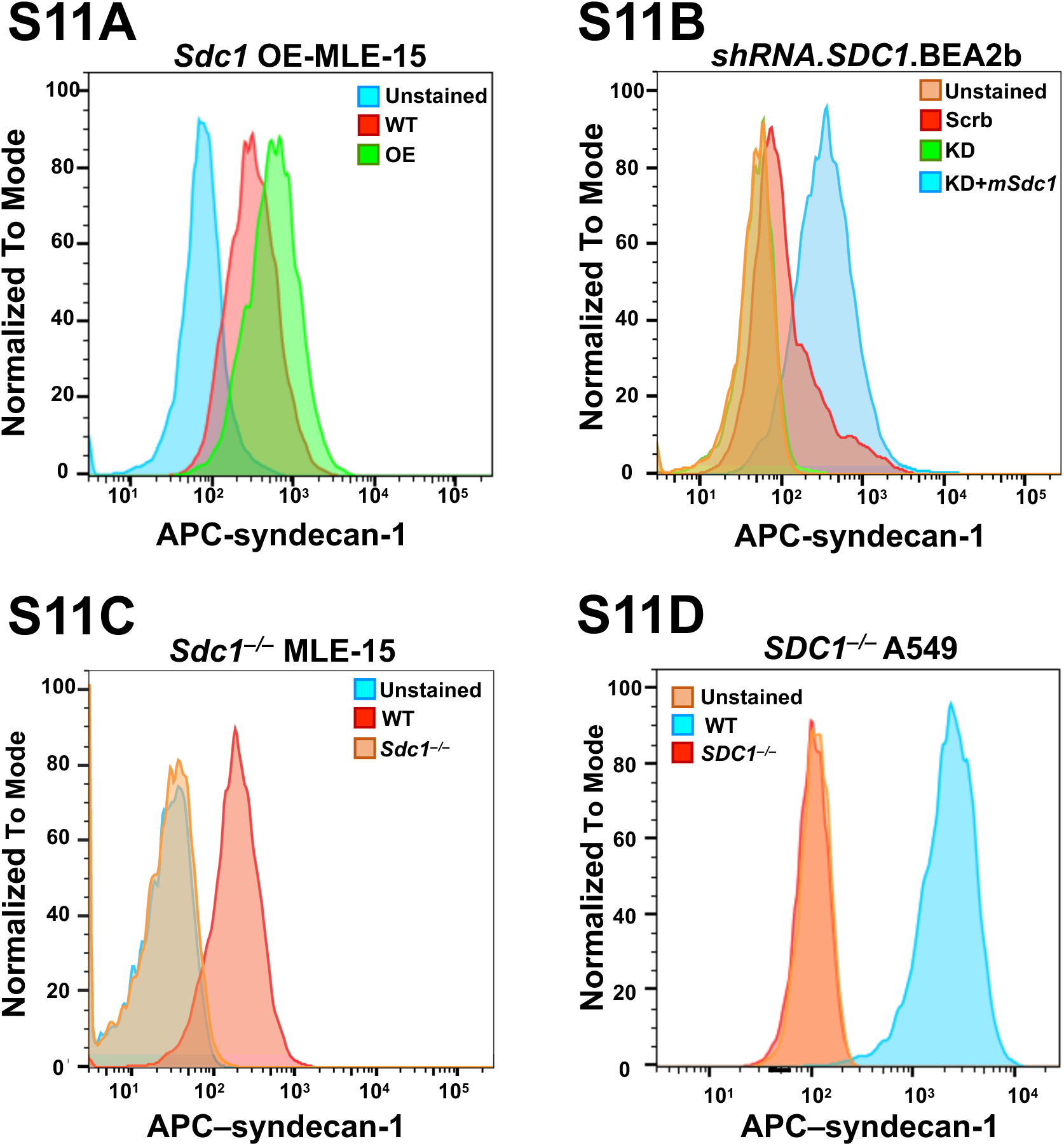
S11A–D Flow cytometric analysis of syndecan-1 (SDC1) expression in mouse and human lung epithelial cell lines. **(A)** *Sdc1* overexpression (OE) in MLE-15 cells. **(B)** *SDC1* silencing and restoration in BEAS-2B cells. **(C)** CRISPR-mediated *Sdc1* deletion in MLE-15 cells. **(D)** CRISPR-mediated *SDC1* deletion in A549 cells.

**Figure S12.**
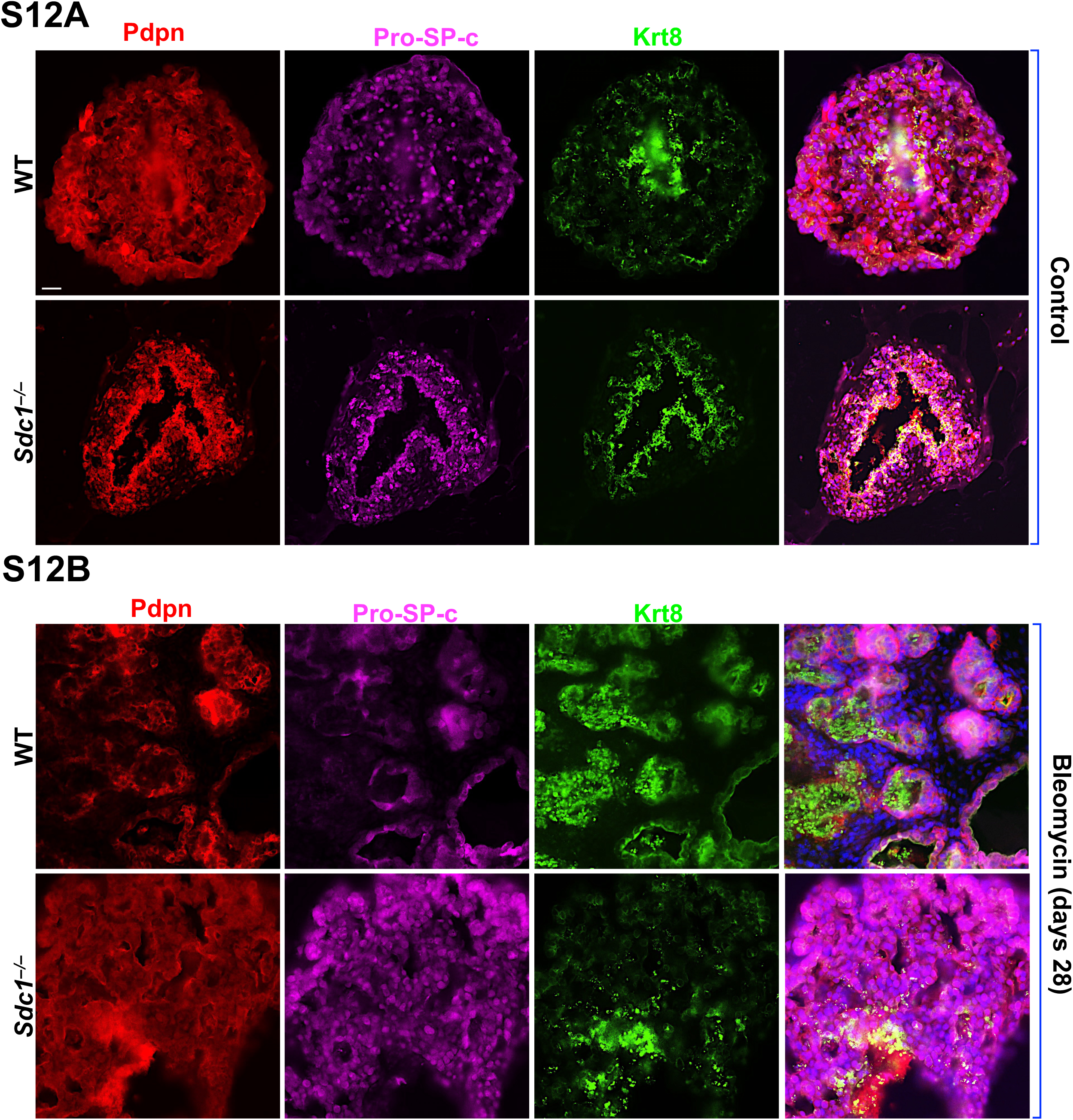
S12A–B Representative immunofluorescence images of AT2 alveolospheres of uninjured (controls, **(A)**) and bleomycin-injured WT and *Sdc1^–/–^* mice **(B)**) stained for AT1 cells (podoplanin [Pdpn]; Alexa Fluor 568, red), AT2 cells (Pro–SP-C; Alexa Fluor 647, magenta), and keratin-8 (Krt8; Alexa Fluor 488, green), N=3 per group. Scale bar=20 µm.

**Figure S13.**
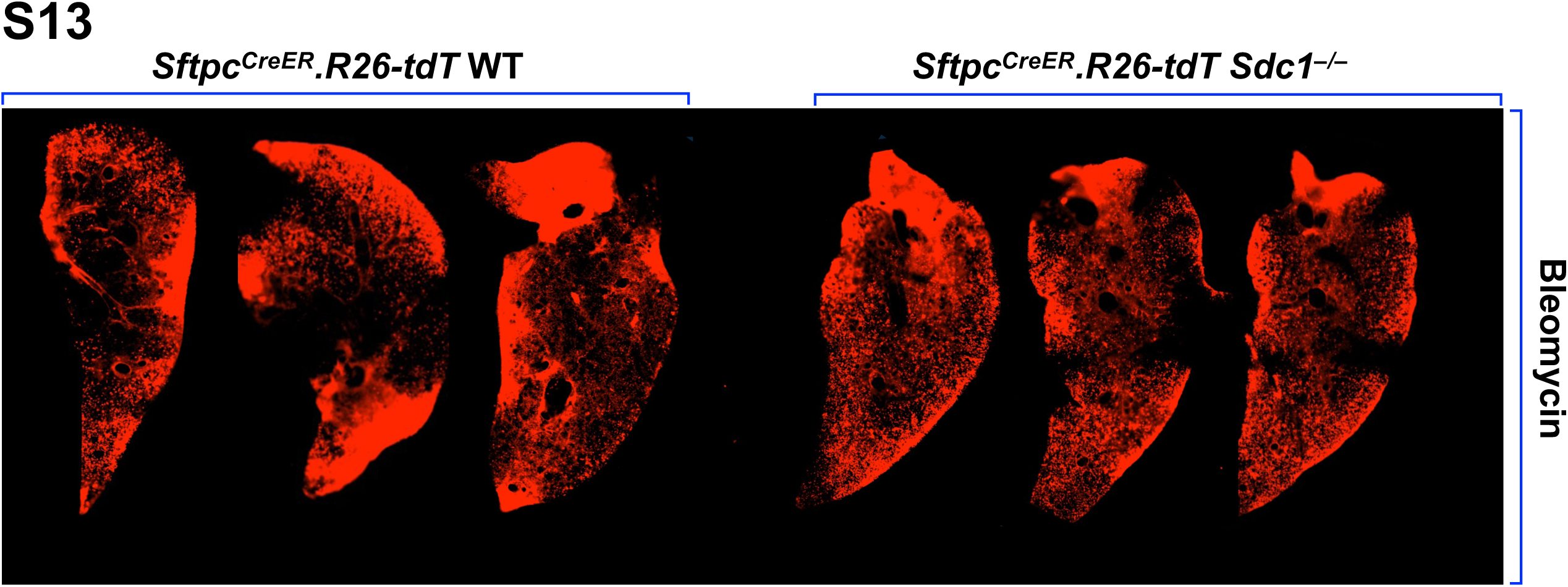
S13 Precision-cut lung slices (PCLS) from young day 14 following bleomycin injury, comparing Sftpc.Rosa26.tdT.CreER.WT and Sftpc.*^Rosa26.tdT.CreER.^ Sdc1^–/–^* genotype, cultured for 7 days and imaged using the Incucyte system. tdTomato (Red) fluorescence reported surfactant protein C-expressing cells, N=3 per group.

**Figure S14.**
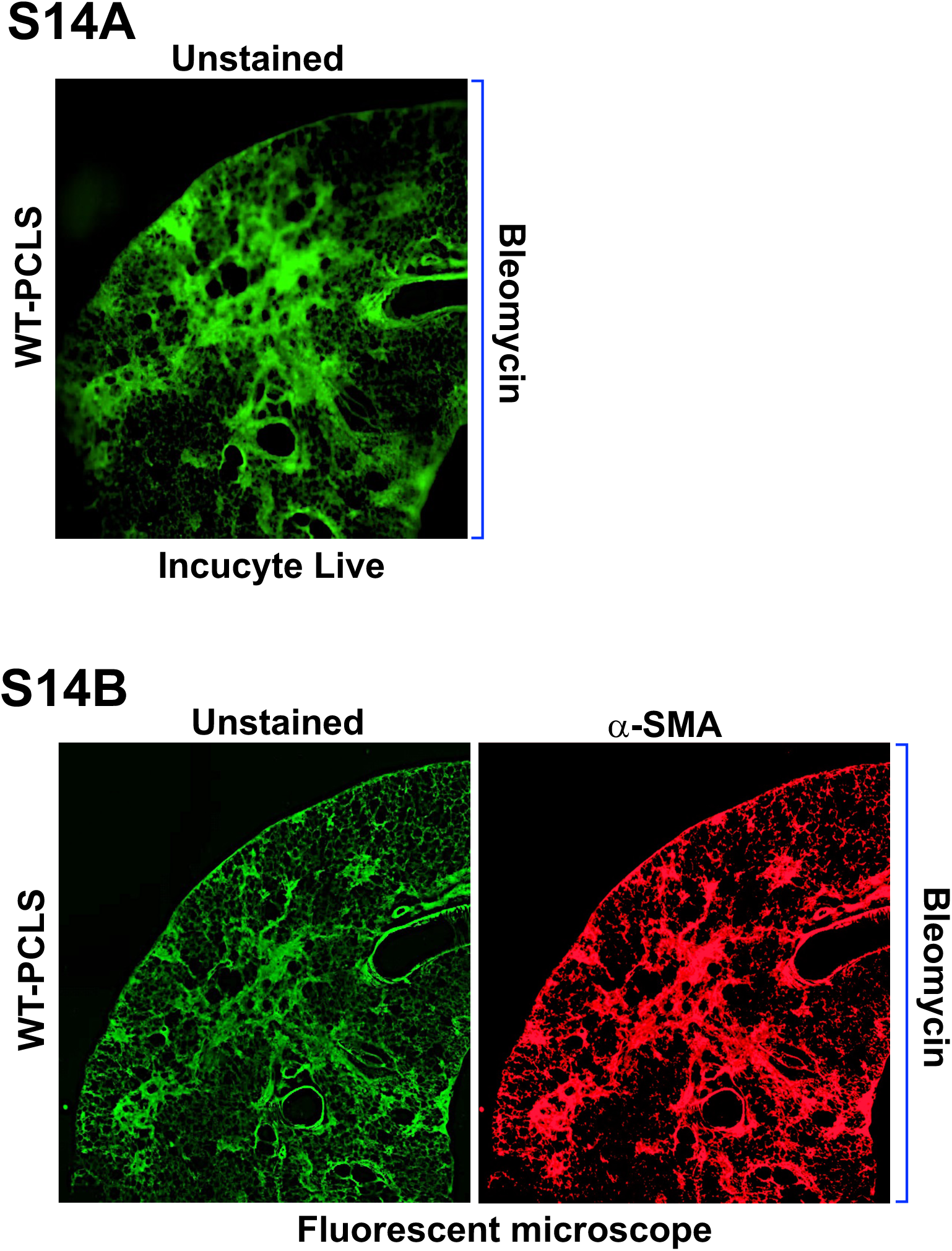
S14A–B Representative images of unstained precision-cut lung slices (PCLS) from bleomycin-injured wild-type (WT) mice. **(A)** acquired by live-cell imaging using the IncuCyte green fluorescent channel and **(B)** a conventional epifluorescence microscope. Corresponding PCLS were subsequently immunofluorescently stained for α–smooth muscle actin (α-SMA; Alexa Fluor 546, Red).

**Figure S15.**
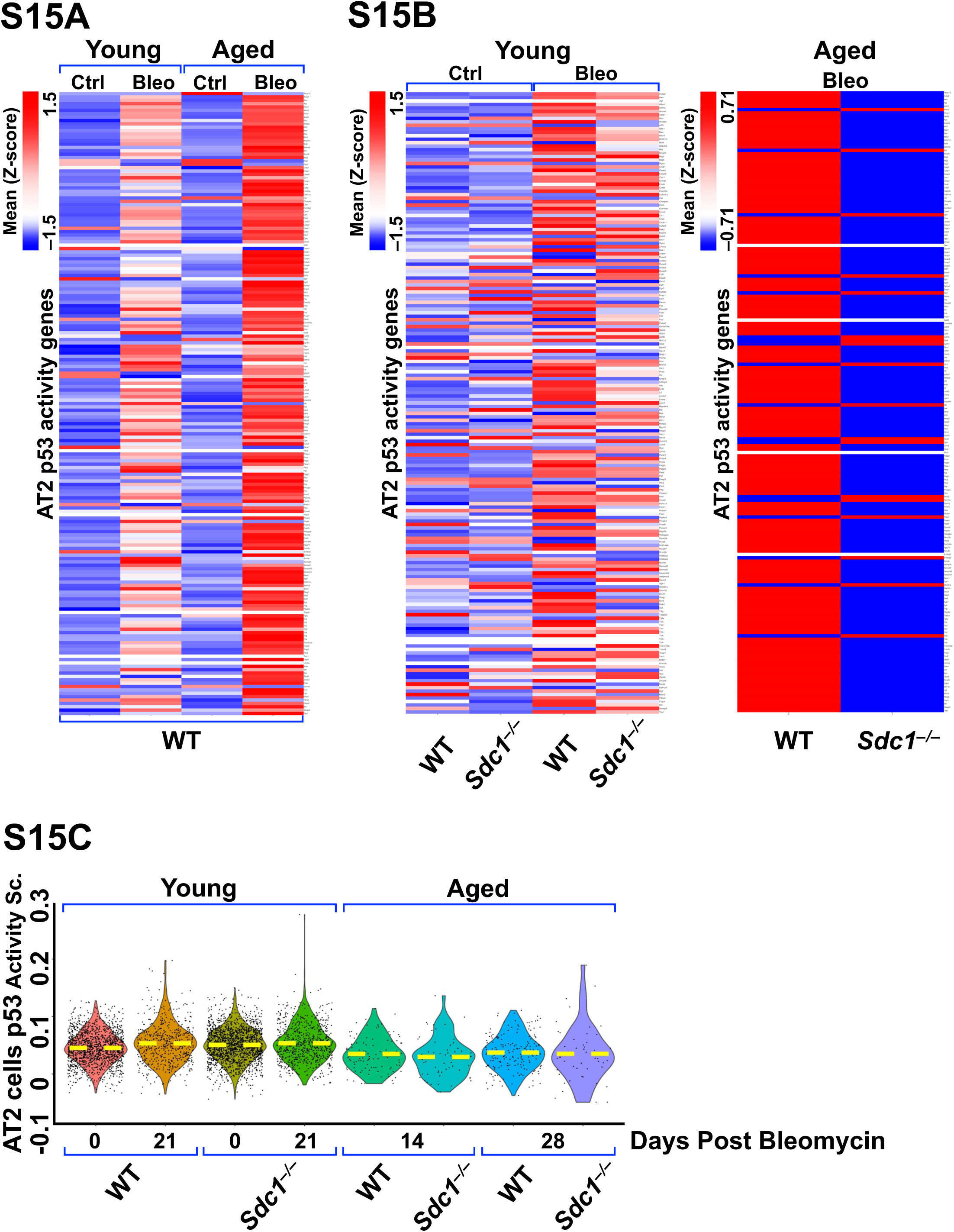
S15A–B Heatmaps illustrating gene expression profiles underlying the p53 activity score in AT2 cells. **(A)** AT2 cells from young and aged WT mice under uninjured and bleomycin-injured conditions (**GSE186246**). **(B)** AT2 cells from young WT and *Sdc1^–/–^* mice following bleomycin injury (**GSE134948**) and from aged bleomycin-injured WT and *Sdc1^–/–^* mice (**GSE292961**). **S15C** Violin plots showing the cumulative mean p53 activity score in AT2 cells, with corresponding heatmaps in **S14B**, from control and bleomycin-injured young WT and *Sdc1^–/–^* mice (**GSE134948**) and aged bleomycin-injured WT and *Sdc1^–/–^* mice (**GSE292961**).

## Unedited Blots

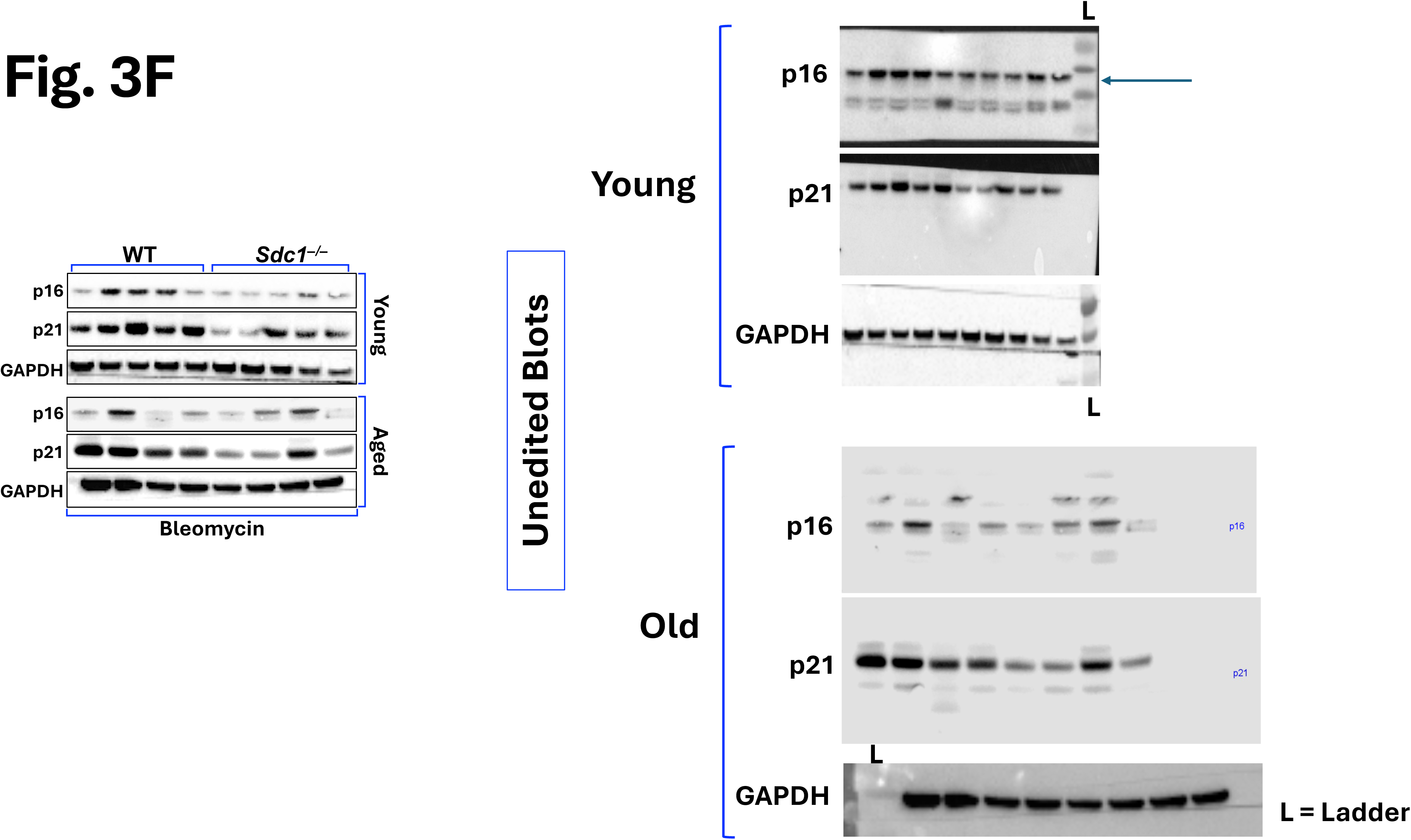

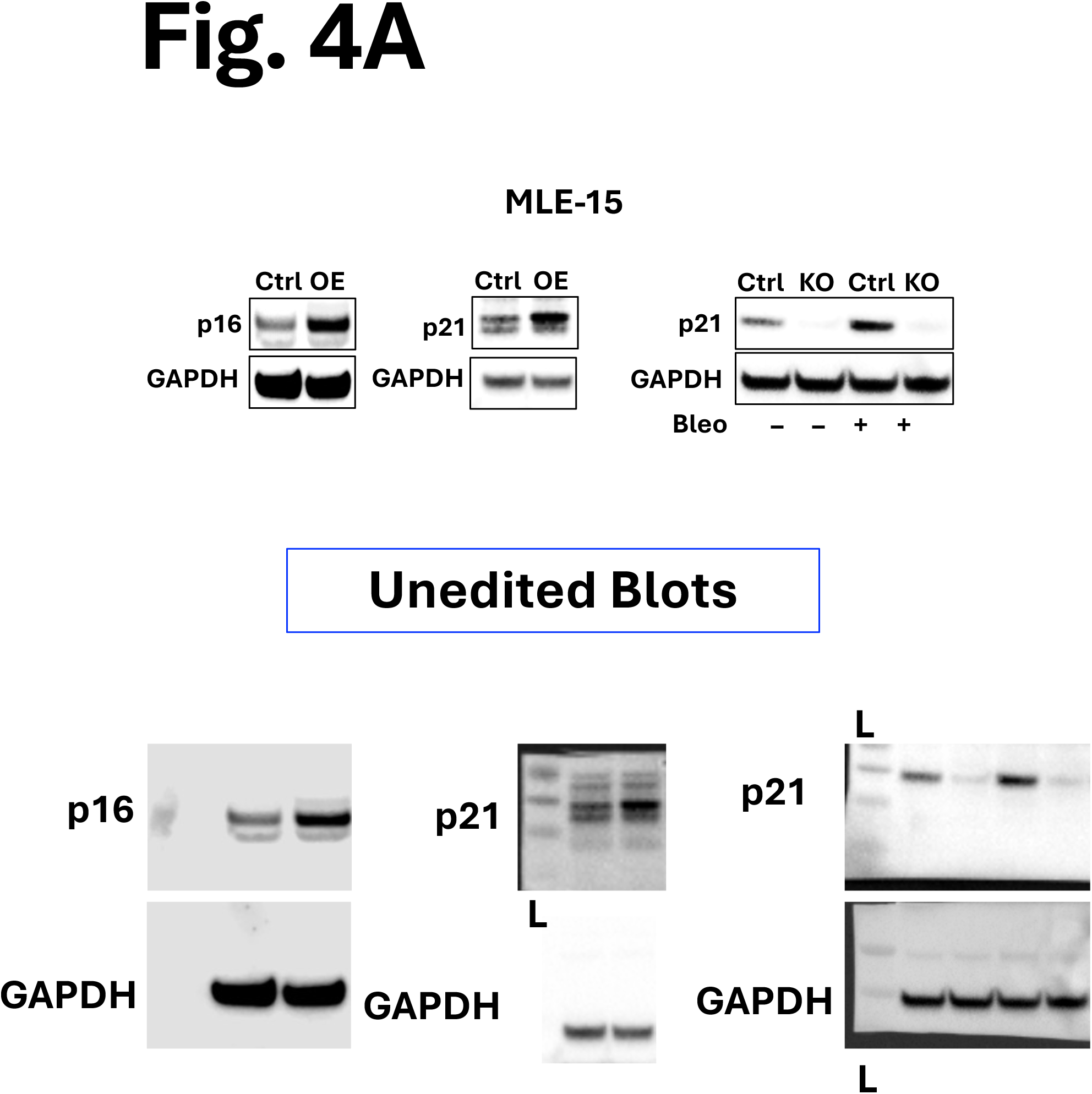

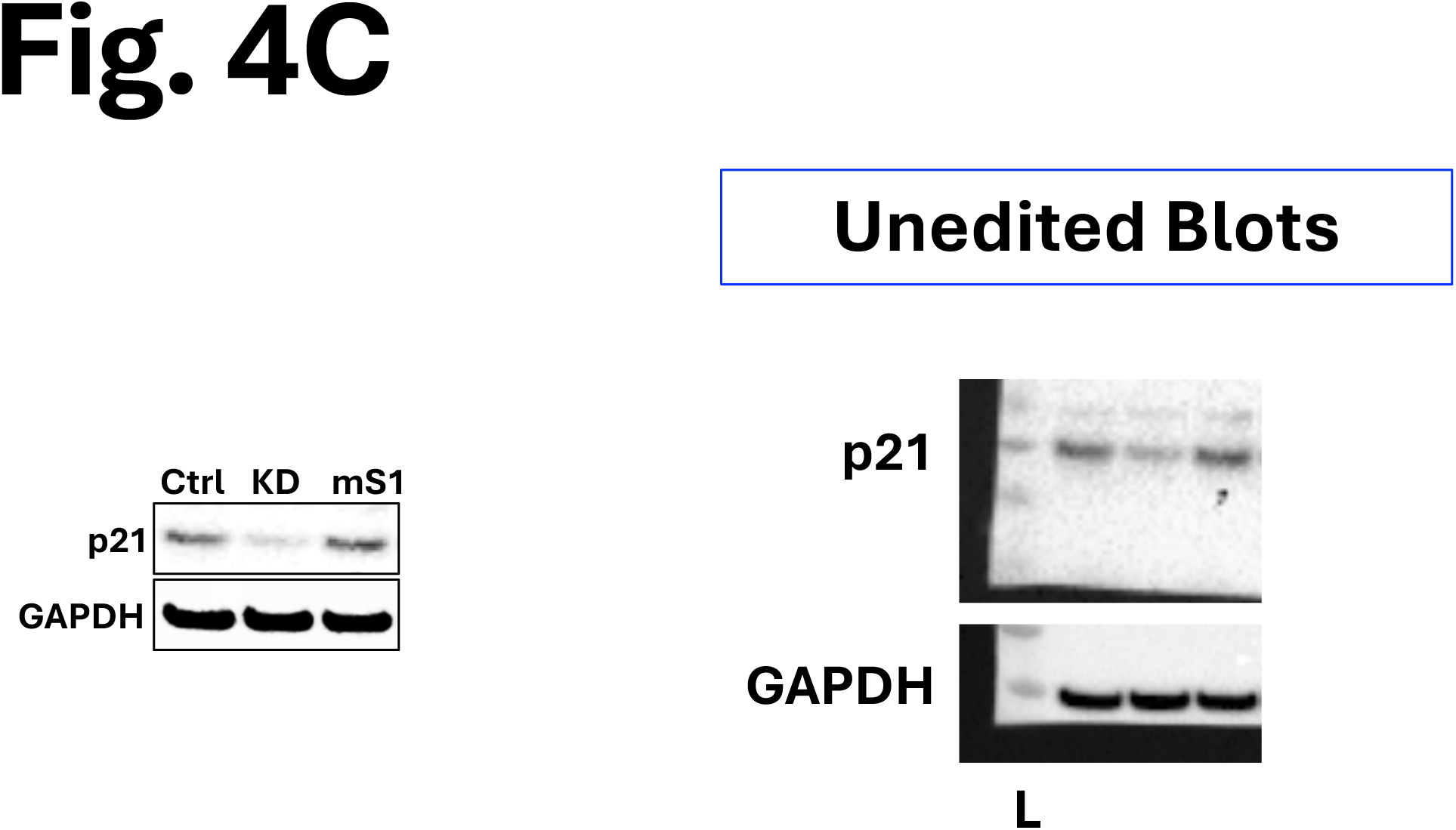

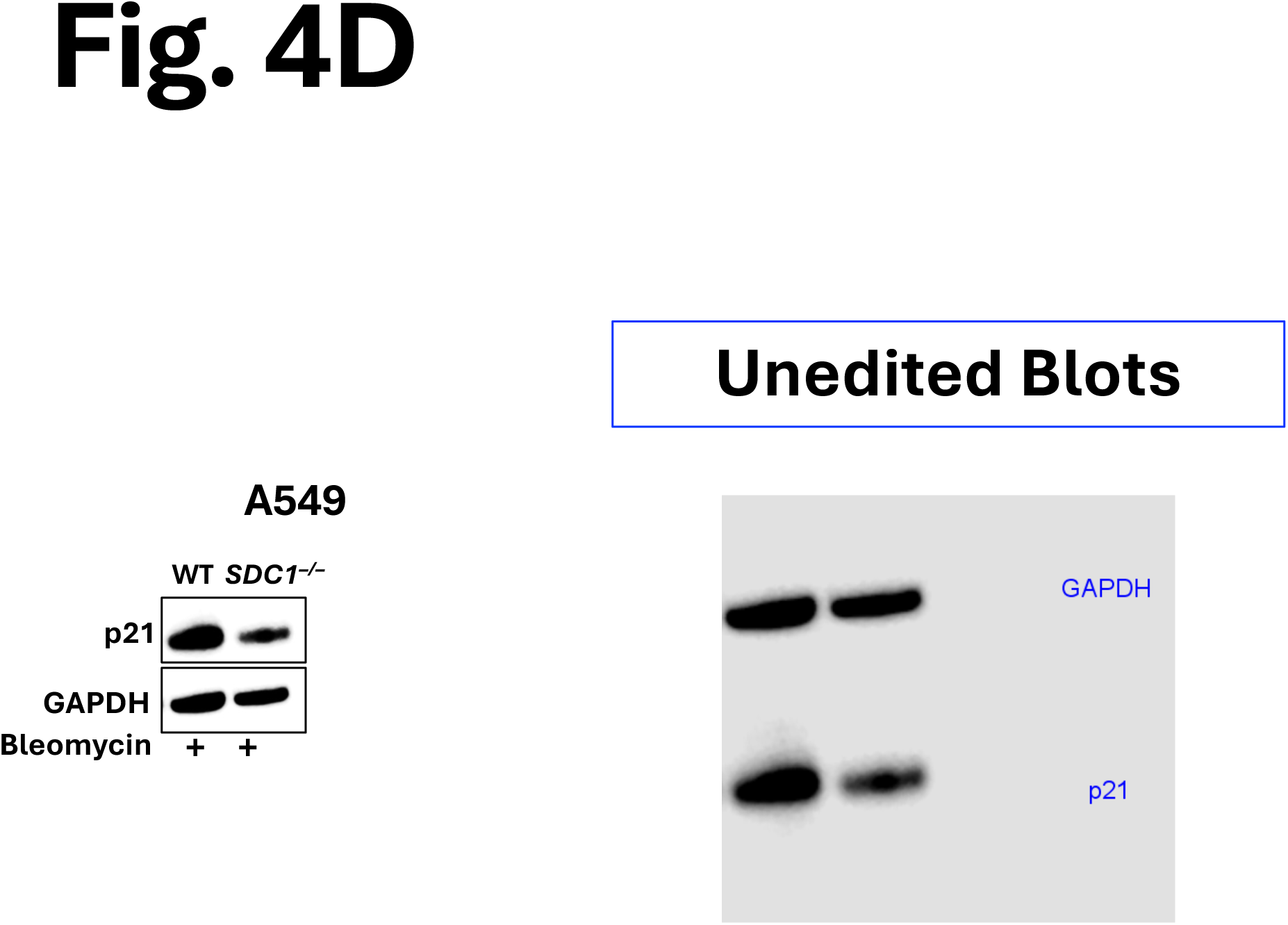

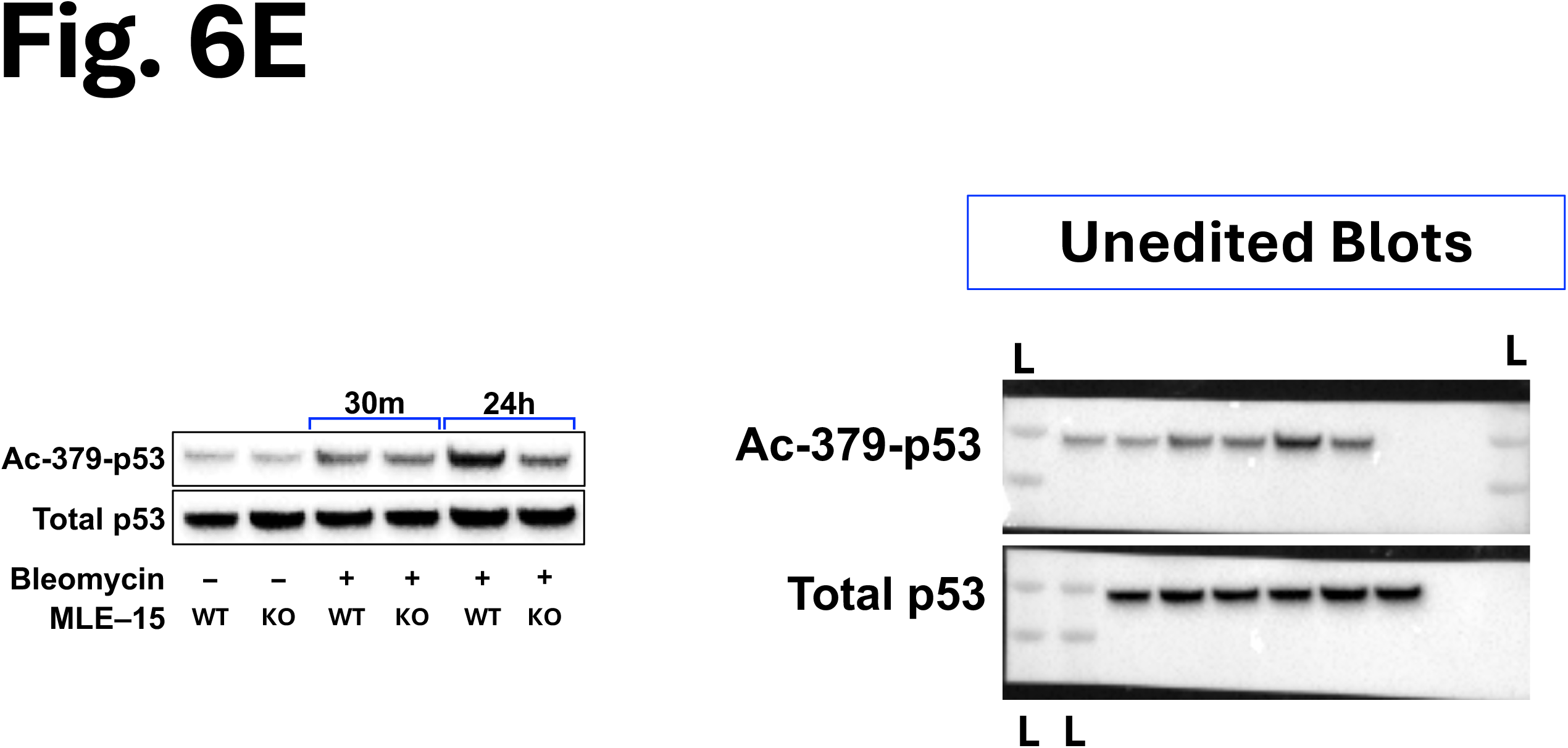

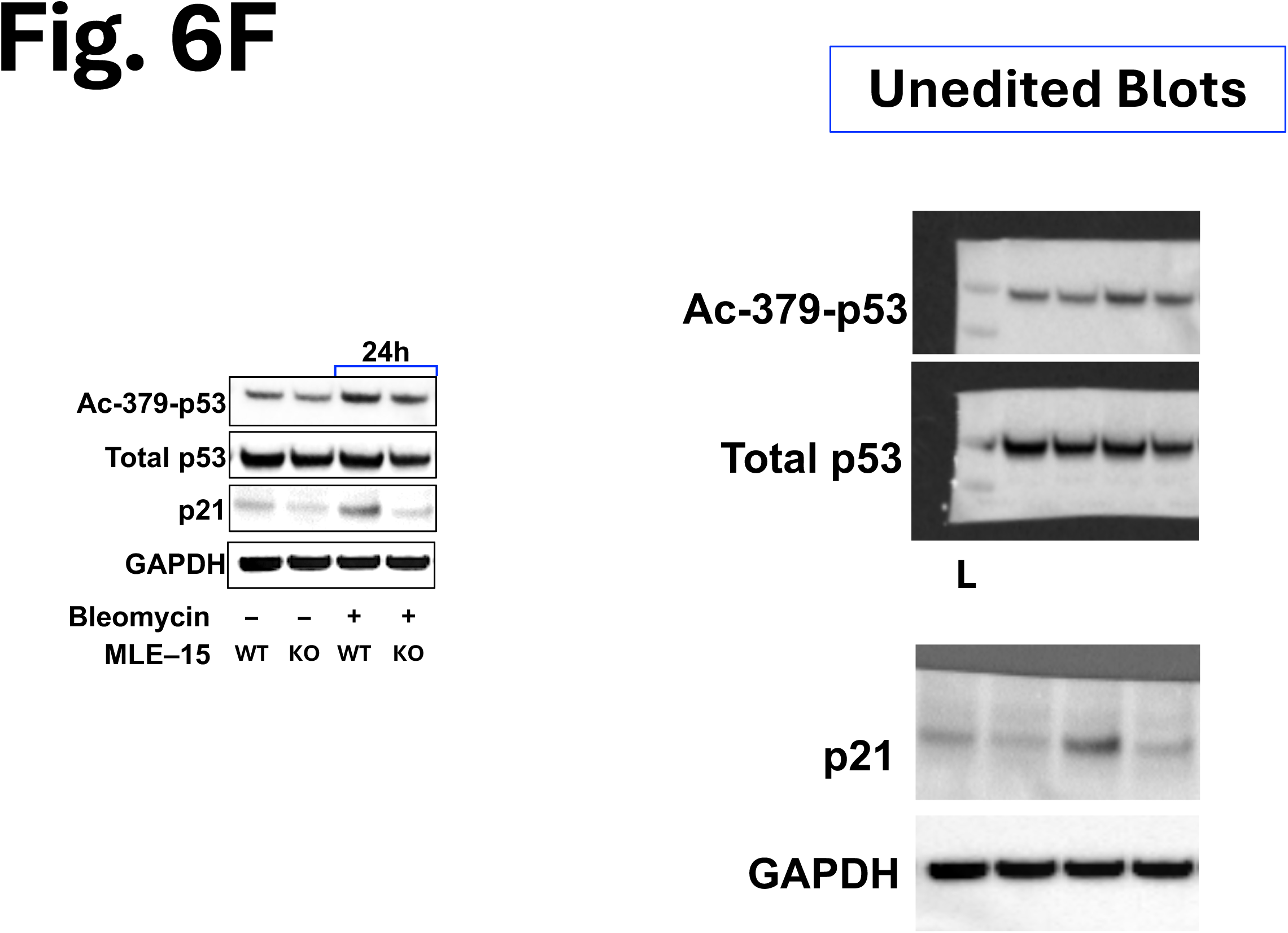

